# *De novo* stem cell establishment in meristems requires repression of organ boundary cell fate

**DOI:** 10.1101/2022.02.03.478990

**Authors:** Antoine Nicolas, Aude Maugarny-Calès, Bernard Adroher, Liudmila Chelysheva, Yu Li, Jasmine Burguet, Anne-Maarit Bågman, Margot E. Smit, Siobhan M. Brady, Yunhai Li, Patrick Laufs

## Abstract

Stem cells play important roles in animal and plant biology as they sustain morphogenesis and tissue replenishment following aging or injuries. In plants, stem cells are embedded in multicellular structures called meristems and the formation of new meristems is essential for the plastic expansion of the highly branched shoot and root systems. In particular, axillary meristems that produce lateral shoots arise from the division of boundary domain cells at the leaf base. The *CUP-SHAPED COTYLEDON* (*CUC*) genes are major determinants of the boundary domain and are required for axillary meristem initiation. However, how axillary meristems get structured and how stem cells become established *de novo* remains elusive. Here, we show that two NGATHA-LIKE transcription factors, DPA4 and SOD7, redundantly repress *CUC* expression in the initiating axillary meristem. Ectopic boundary fate leads to abnormal growth and organisation of the axillary meristem and prevents *de novo* stem cell establishment. Floral meristems of the *dpa4 sod7* double mutant show a similar delay in stem cell *de novo* establishment. Altogether, while boundary fate is required for the initiation of axillary meristems, our work reveals how it is later repressed to allow proper meristem establishment and *de novo* stem cell niche formation.

## INTRODUCTION

Stem cells play a central role in animal and plant biology as they are the source of all cells that form organs and tissues during morphogenesis and allow cells to be replaced following injuries or at the end of their life cycle (Baurle and Laux, 2003; Birnbaum and Alvarado, 2008; Morrison and Spradling, 2008). In both animals and plants, stem cells are maintained in their undifferentiated and pluripotent state through interactions with a microenvironment that forms a niche (Comazzetto et al., 2021; Dinneny and Benfey, 2008; Janocha and Lohmann, 2018; Pardal and Heidstra, 2021; Xie and Spradling, 2000). However, in contrast to what occurs in animals, plant stem cells cannot move and, as a consequence, stem cell niches have to be formed *de novo* in plants (Laird et al., 2008). Indeed, *de novo* stem cell establishment is essential to support the formation of new growth axes (shoots or roots) that allows plants to plastically expand their shape enabling them to explore their environment.

In plants, stem cells and niches are embedded in multicellular structures called meristems. The shoot apical meristem (SAM), formed during embryogenesis, is the direct source of the main shoot, forming stem and leaves after germination (Long et al., 1996). The SAM is a dynamic, yet organized structure that is maintained through interactions between its different domains. In the apical part of the SAM lies a group of semi-permanent stem cells maintained by an underlying organizing centre (OC) that contributes to the stem cell niche function (Laux et al., 1996). The OC expresses the WUSCHEL (WUS) transcription factor that travels through cellular connections to the overlying layers to induce stem cell fate (Daum et al., 2014; Mayer et al., 1998; Perales et al., 2016; Sloan et al., 2020; Yadav et al., 2011). In turn, stem cells express the excreted CLAVATA3 (CLV3) peptide that through interaction with different receptor kinases including CLAVATA1 (CLV1) feedbacks to repress *WUS* activity in the OC (Brand et al., 2000; Fletcher et al., 1999; Müller et al., 2008; Schlegel et al., 2021; Schoof et al., 2000). On this core *WUS*/*CLV* regulatory feedback circuit are grafted additional interacting regulators such as auxin and cytokinin signals or the HAIRY MERISTEM (HAM) transcription factors and their regulatory miRNA, miR171 (Chickarmane et al., 2012; Gruel et al., 2016; Han et al., 2020a; Leibfried et al., 2005; Ma et al., 2019; Zhou et al., 2015). Altogether, this network contributes to proper spatial positioning of the stem cell and stem cell niche and their fine tuning to allow meristem activity to respond to environmental signals (Landrein et al., 2018; Pfeiffer et al., 2016; Yoshida et al., 2011).

On the flanks of the meristem, new organ primordia are initiated following a spatial and temporal pattern that is orchestrated by auxin and cytokinin signaling (Besnard et al., 2014; Reinhardt et al., 2003). Proper initiation and separation of the organ primordia requires the establishment of an organ boundary domain by multiple factors in which the *CUP-SHAPED COTYLEDON* (*CUC*) genes play a prominent role (Aida and Tasaka, 2006; Žádníková et al., 2014). This domain separates the leaf primordium from the meristem and will later give rise to the axillary region that lies on the inner base of the leaf. Multiple factors allow coordinating primordium initiation with stem cell activities. For instance, the *CUC* genes are both required for organ formation and meristem maintenance (Aida et al., 1997), the HD-ZIP III transcription factors contribute to leaf polarity and meristem function (Caggiano et al., 2017; Kim et al., 2008), and auxin and cytokinins are regulating both organ initiation and stem cell activity (Besnard et al., 2014; Chickarmane et al., 2012; Ma et al., 2019; Reinhardt et al., 2003).

While the root and shoot apical meristems formed during embryogenesis are generating respectively the primary root and the main shoot, the ramified architectures of the shoot and root systems result from the activity of meristems newly formed during post-embryonic development. These lateral root and shoot meristems arise from a group of dividing cells originating respectively from the root pericycle layer or the leaf axillary region and acquire an organization and activity similar to the primary embryonic meristems, including a *de novo* established stem cell population and niche.

The formation of an axillary meristem (AM) between the developing leaf primordia and the SAM can be divided into three steps: the maintenance of a few meristematic cells at the leaf axil, the expansion of this cell population and the establishment of a functional meristem (Cao and Jiao, 2020; Wang and Jiao, 2018; Wang et al., 2016). Multiple factors regulating these events have been characterized during the formation of the AM formed in the rosette leaves of *Arabidopsis thaliana*. During the maintenance phase, a small group of cells located at the base of the developing leaf retains meristematic features while neighboring cells differentiate (Grbic and Bleecker, 2000; Long and Barton, 2000). Expression of the meristematic gene *SHOOT MERISTEMLESS* (*STM*) and the boundary domain genes *CUC2* and *CUC3* in these cells is required for AM initiation and, accordingly, *stm*, *cuc2* or *cuc3* mutants show defective AM formation (Grbic and Bleecker, 2000; Hibara et al., 2006; Long and Barton, 2000; Raman et al., 2008; Shi et al., 2016). Maintenance of *STM* expression requires auxin depletion from the axillary region by polar auxin transport (Wang et al., 2014a, 2014b) and involves at the molecular level a self-activation loop facilitated by a permissive epigenetic environment (Cao et al., 2020). These cells can remain latent during a long period of time and, upon receiving proper environmental or endogenous signals, switch to the activation phase during which their number rapidly increases by cell divisions to generate a small bulge. A strong increase in *STM* expression level is instrumental for the switch to the activation phase (Shi et al., 2016) and multiple transcription factors, such as REVOLUTA, DORNRÖSCHEN, DORNRÖSCHEN LIKE, REGULATOR OF AXILLARY MERISTEMS1, 2 and 3 and REGULATOR OF AXILLARY MERISTEM FORMATION provide spatial and temporal cues for the local activation of *STM* expression (Greb et al., 2003; Keller et al., 2006; Müller et al., 2006; Raman et al., 2008; Shi et al., 2016; Yang et al., 2012; Zhang et al., 2018). Furthermore, a local pulse of cytokinin signalling reinforces *STM* expression to promote the formation of the AM, possibly through a mutual positive feedback loop between STM and cytokinins (Wang et al., 2014b).

During the establishment phase, the bulge acquires progressively a typical meristem organization with functional sub-domains. Cytokinins promote *de novo WUS* expression, thus defining the OC (Wang et al., 2017). For this, the type-B Arabidopsis response regulator proteins (ARRs), which mediate the transcriptional response to cytokinin, directly bind to the WUS promoter. In turn, *WUS* expression initiates the activation of the stem cell population marked by the expression of the *CLV3* gene (Xin et al., 2017). Interestingly, during the initial phase of *WUS* and *CLV3* activation both genes are expressed in overlapping domains in internal layers of the AM and they only later discriminate into their proper expression patterns with *CLV3* expression shifting to the upper layers (Xin et al., 2017). This spatial rearrangement of *CLV3* expression requires an apical-basal gradient of *HAM* genes activities that results in part from the epidermis-specific expression of their negative regulators miR171 (Han et al., 2020a, 2020b; Zhou et al., 2018). Therefore, AM establishment is a gradual process, during which the expanding population of meristematic cells acquires specific identities including the specification of apical stem cells and an underlying stem cell niche combined with organ boundary domains at the meristem flanks.

Arabidopsis floral meristems are proposed to be modified AMs in which the subtending leaf is replaced by a cryptic bract whose development is suppressed (Long and Barton, 2000). Floral meristems also establish *de novo* a stem cell population marked by a rapid activation of *WUS* expression in stage 1 floral meristems (Mayer et al., 1998) and by *CLV3* expression by late stage 2 (Seeliger et al., 2016). However, in contrast to AM, in which stem cells are maintained, floral meristems are determined structures with only a transient maintenance of stem cells. Indeed, the C-class floral gene *AGAMOUS* directly repress *WUS* by recruiting the Polycomb Repressive Complex 1 factor TERMINAL FLOWER 2 and induces KNUCKLES which in turn represses *WUS* and interferes with WUS-mediated CLV3 activation (Lenhard et al., 2001; Liu et al., 2011; Shang et al., 2021; Sun et al., 2014).

Thus, it appears that while the molecular mechanisms allowing the preservation and the amplification of a pool of meristematic cells leading to AM emergence start to be deciphered, how the newly meristem becomes organized and activated remains far less understood. Here, we analyse AM establishment, concentrating on cauline AMs (CaAMs) that are poorly characterized compared to rosette AMs (RoAMs). We show that CaAMs are rapidly formed following floral induction and that this is associated with dynamic changes in gene expression. Accordingly, while the *CUC* genes are required for the maintenance and activation phase, they have to be cleared for meristem establishment and activation of the stem cell population. Indeed, ectopic expression of the *CUC* boundary genes leads to asynchronous AM development and delayed *de novo* stem cell formation. We provide a molecular mechanism for this dynamic regulation of the *CUC* genes by two members of the NGATHA-like (NGAL) family of transcriptional repressors. A similar delay in *de novo* stem cell establishment is observed during floral meristem formation. Altogether, we reveal a genetic circuit repressing boundary cell fate that is required for *de novo* stem cell formation.

## RESULTS

### Dynamic gene expression accompanies cauline AM establishment

CaAMs are rapidly formed and grow out following floral transition (Burian et al., 2016; Grbic and Bleecker, 2000; Hempel and Feldman, 1994). To provide a framework for CaAM formation in Arabidopsis, we analysed morphological changes in calcofluor-stained samples and gene expression dynamics using reporter lines in the leaf axillary region following plant shifting from short-day (SD) to long-day (LD) conditions (Fig. 1). Six days after shifting to LD (6LD), the cauline leaf primordium was separated from the main meristem by a boundary containing small and narrow cells (Fig. 1A). At 8LD, a bulge emerged between the cauline leaf primordium and the i, defining the “dome stage” of the developing AM (Fig. 1B). At 10LD and 13LD leaf and flower primordia were formed by the AM (Fig. 1C,D), defining respectively the “leaf primordium” and “flower primordium” stages.

**Figure 1.**
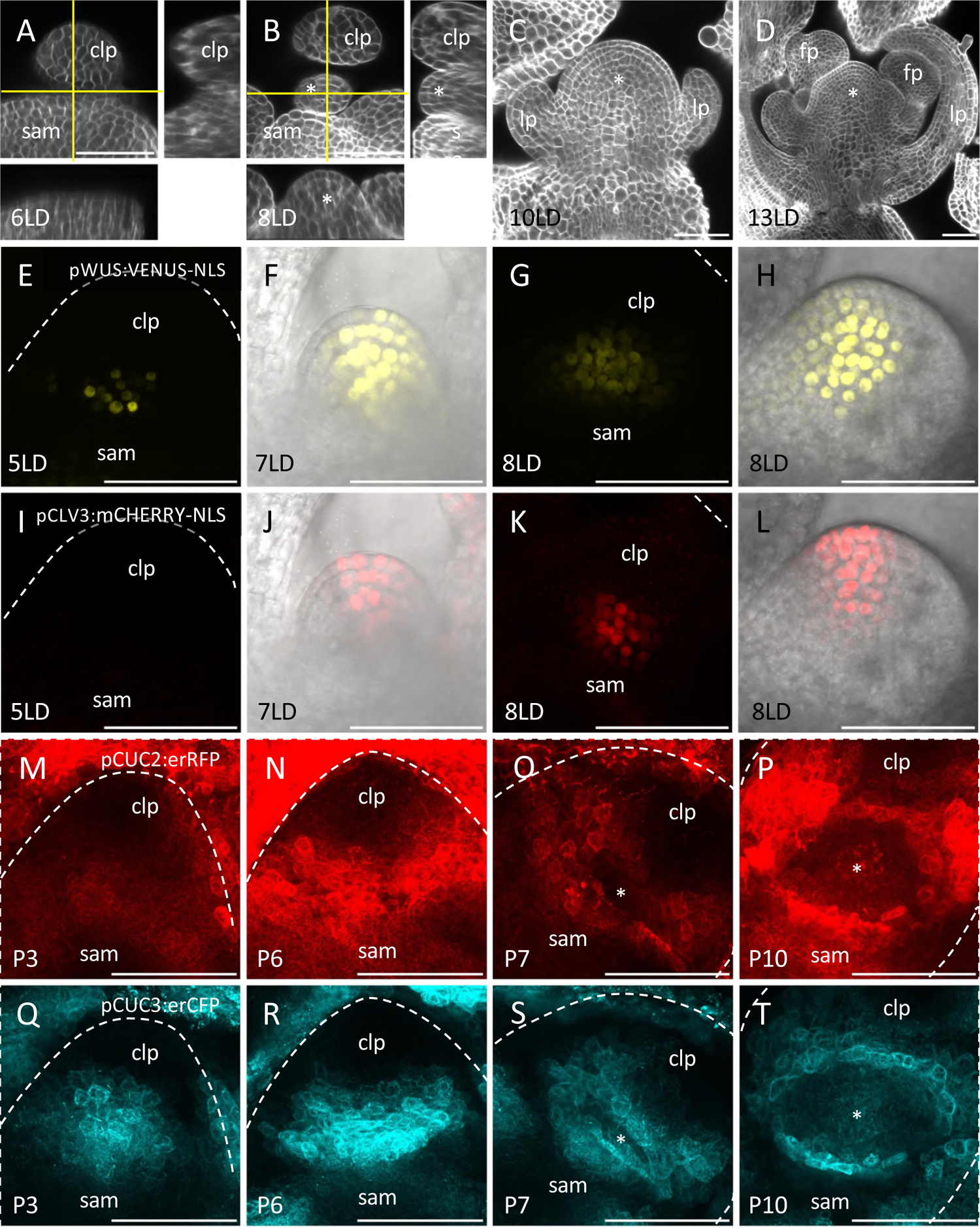
Rapid morphological changes and dynamic gene expression accompany CaAM formation. (A-D) Optical sections of calcofluor-stained axillary regions of wild-type following SD to LD transition. (A-B) Main panel: transverse optical section (with respect to the main stem axis), lower panel: reconstructed optical tangential section, right panel: reconstructed optical radial sections. Yellow lines mark the position of the tangential and radial sections. (C-D) Optical tangential sections. The number of days in LD condition is indicated. (E-L) Maximum projections of transverse (E,G,I,K) and tangential (F,H,J,L) optical sections of a pWUS:VENUS-NLS (E-H) and pCLV3:mCHERRY-NLS (I-L) reporter line during CaAM formation. (F,J,H and L) are a merge between reporter fluorescence and transmitted light. The number of days in LD condition is indicated. (M-T) Maximum projections of transverse optical sections of a pCUC2:erRFP (M-P) and pCUC3:erCFP (Q-T) reporter line during CaAM formation. Positions are numbered according to the rank of the primordium. Primordium number is indicated. Scale bars = 50µm; sam: shoot apical meristem; clp: cauline leaf primordium; *: AM; lp: leaf primordium formed by the AM, fp: flower primordium formed by the AM. The dotted line corresponds to the outline of the cauline leaf primordium.

To trace back the formation of the organizing centre and stem cells during CaAM establishment, we first analysed the expression dynamics of *WUSCHEL* and *CLAVATA3* transcriptional reporters (Pfeiffer et al., 2016). At 5LD, pWUS:3xVENUS-NLS expression appeared in a few cells in of P7, the 7^th^ youngest visible primordia (Fig. 1E). The number of VENUS expressing cells progressively increased during later stages (Fig. 1F-H). pCLV3:mCHERRY-NLS expression appeared only later: some CaAMs started to express *CLV3* at 7LD while at 8 LD most of them expressed *CLV3* (Fig.1 I-L). Longitudinal optical sections showed that at 7LD *WUS* expression expanded from the corpus into the L2 and sometimes L1 layer (Fig. 1F). Concomitant with the onset of *CLV3* expression (Fig. 1L), *WUS* expression became progressively excluded from the 3 outermost layers to finally mimic the expression observed in the SAM (Fig. 1H). Therefore, as in the RoAMs, during *de novo* establishment of the stem cell niche in CaAM, *WUS* is first activated, while *CLV3* is expressed later in a domain contained in the WUS-expressing cells. These two overlapping domains then resolve into an apical *CLV3* domain and a central *WUS* domain. However, whereas in the RoAMs, *CLV3* showed a dynamic shift from a central to an apical domain (Xin et al., 2017) (Fig. S1), in CaAMs, the WUS domain shifted from an apical to a central domain.

Next, we followed the dynamics of *CUC2* and *CUC3* expression as these genes are redundantly required for CaAM formation (Hibara et al., 2006; Raman et al., 2008). During the early stages (P1 to P6), pCUC2:erRFP and pCUC3er:CFP transcriptional reporters showed a compact domain of expression at the boundary between the cauline leaf primordia and the meristem (Fig. 1M,Q). These expression domains became progressively more elongated while the groove separating the primordium from the meristem formed (P5-P6, Fig. 1N,R). Such *CUC2* and *CUC3* expression dynamics were independent of CaAM formation as they were observed in apices of both plants shifted or not shifted to LD. However, while in non-induced plants, expression of *CUC2* and *CUC3* remained as a compact line, we observed that starting 5LD onwards, it split into an eye-shaped structure leaving a central region with reduced expression in P7-P8 primordia (Fig. 1O,S). The central domain depleted for *CUC2* and *CUC3* expression expanded during later stages (P9-P10 in plants at >6LD), the expression of the two reporters concentrating into a necklace-shaped structure around the outgrowing meristematic dome (Fig. 1P,T).

In conclusion, CaAM formation is a rapid process leading to *de novo* establishment of a novel functional meristem, containing an organizing center and stem cell population. *CUC2* and *CUC3* expression is dynamic during CaAM formation shifting from an expression throughout the meristem to an expression restricted around the meristem.

### Identification of putative regulators of *CUC* gene dynamic expression

To identify possible transcriptional regulators of the dynamic expression of the *CUC2* and *CUC3* genes during AM formation, we performed an enhanced yeast one-hybrid screen using the *CUC2* and *CUC3* promoter regions as baits (Gaudinier et al., 2011). Thus, we identified SOD7/NGAL2 as a protein binding to the *CUC3* promoter. SOD7/NGAL2 is a member of the small family of NGATHA-like transcription factors (Romanel et al., 2009; Swaminathan et al., 2008). We did not detect any interaction with ABS2/NGAL1, while DPA4/NGAL3 was not present in the transcription factor collection we screened (See Supplementary Material). However, because the *NGAL* genes were shown to repress *CUC* genes during leaf and seedling development (Engelhorn et al., 2012; Shao et al., 2020), we next tested whether the *NGAL* genes could be involved in AM development.

### The *SOD7/NGAL2* and *DPA4/NGAL3* genes are redundantly required for AM formation

To determine if the *NGAL* genes had a role in AM development, we grew single and multiple *ngal* mutants for 5 weeks in LD conditions. Undeveloped or delayed CaAMs were frequently observed in the *dpa4-2 sod7-2* double mutant and *abs1 dpa4-2 sod7-2* triple mutant, compared to the CaAMs in WT and other single or double mutants (Fig. 2A-H). To quantify this phenotype more precisely, we performed a kinetics of CaAM development and calculated the time point after bolting at which half of the CaAMs were developed (t_50_). We observed a delay in the development of CaAMs for the double *dpa4-2 sod7-2* (t_50_ = 6.9 days) and triple *abs1 dpa4-2 sod7-2* (t_50_ = 6.6 days) mutants compared to the WT and the other mutants (t_50_ = 1.5 days) (Fig. 2I, Fig. S2A). A delay in RoAM development was also observed for the *dpa4-2 sod7-2* and *abs1 dpa4-2 sod7-2* mutants (Fig. S2B-D). Finally, the double mutant *dpa4-3 sod7-2* with another *dpa4* mutant allele also showed a delayed CaAM development (Fig. S2G,H). All together, these data show that the *NGAL* genes are redundantly required for CaAM and RoAM development and that *DPA4* and *SOD7* play a major role in this process, while *ABS2* has only a minor contribution.

**Figure 2.**
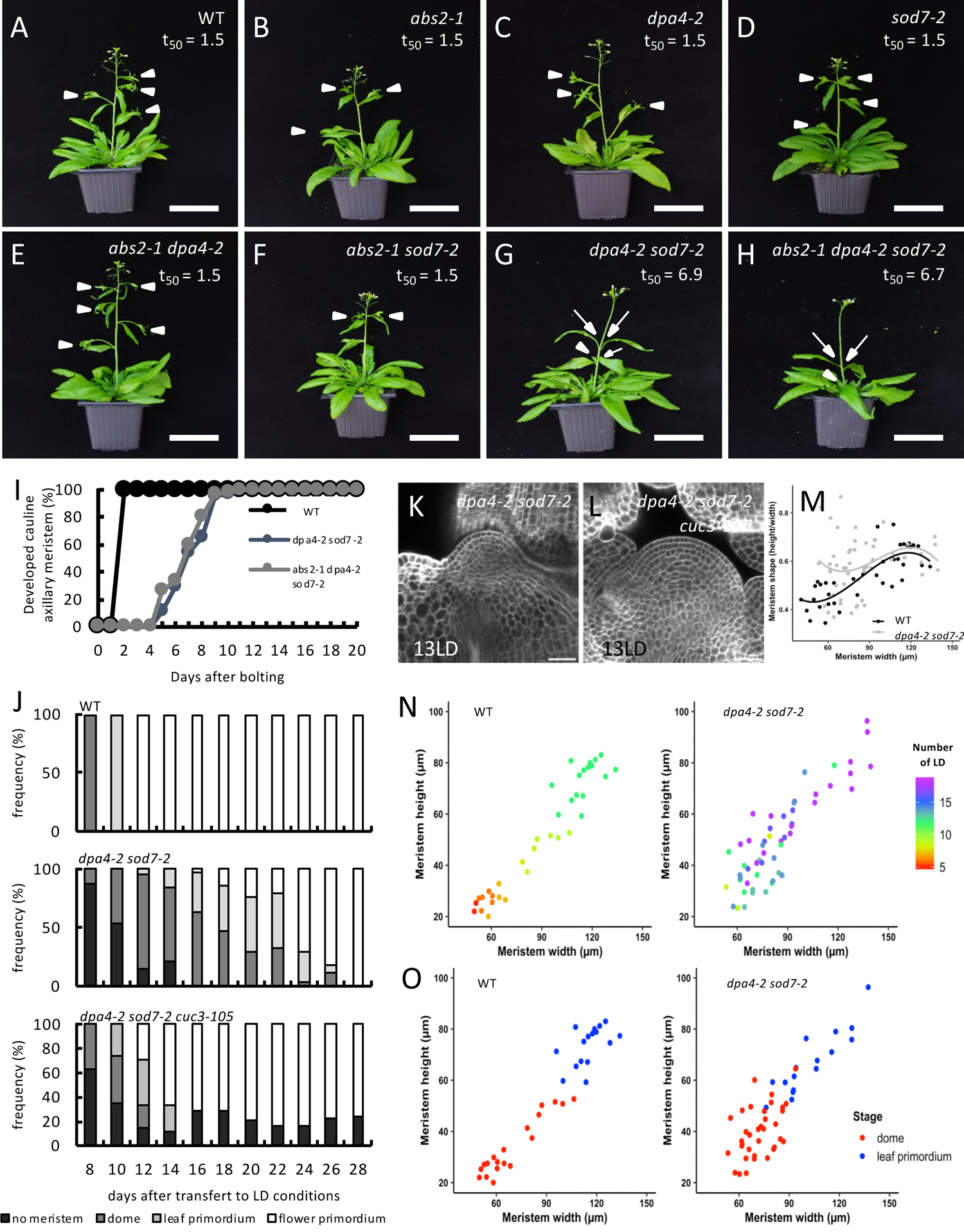
*DPA4* and *SOD7* are required for rapid development of cauline AMs. (A–H) Inflorescence of WT, simple, double and triple *ngal* mutants. Plants were grown for 5 weeks in long-day conditions. White arrowheads point to developed CaAMs while the arrows point to delayed CaAMs. The time point after bolting at which half of the CaAM are developed (t_50_, in days) is indicated under the genotype. (I) Kinetics of CaAM development after bolting. Development of the CaAM is indicated as the percentage of developed branches (≥ 3mm) reported to the total number of cauline leaves (*n*≥11). (J) Kinetics of CaAM development in WT, *dpa4-2 sod7-2* and *dpa4-2 sod7-2 cuc3-105* grown 4 weeks in SD and transferred to LD (*n* ≥ 10). (K,L) Tangential optical sections of calcofluor-stained WT of *dpa4-2 sod7-2* (K) and *dpa4-2 sod7-2 cuc3-105* (L) CaAM at 13LD. The wild-type control is shown in Fig1D (M) Evolution of CaAM shape in WT and *dpa4-2 sod7-2*. (N) CaAM height and width as a function of the number of LD in WT and *dpa4-2 sod7-2*. (O) CaAM height and width as a function of the CaAM stage in WT and *dpa4-2 sod7-2*. Scale bars: (A–H) = 5 cm; (K,L) = 100 µm

Next, we traced back the origin of the delayed AM development by looking at early stages of CaAMs and RoAMs in the WT and the *dpa4-2 sod7-2* mutant. In the WT, all the CaAMs rapidly switched from the dome stage at 8LD, to the leaf primordium stage at 10LD and at the flower primodium stage at 12LD (Fig. 2J, top plot). In contrast, no meristem was visible in the majority of the *dpa4-2 sod7-2* cauline leaves at 8LD, while meristems at the dome stage were present only in about half of the axils at 10LD (Fig. 2J, middle plot). The apparition of leaf primordium and flower primordium was also delayed compared to the WT. In addition, confocal observations of dpa4-2 *sod7-2* meristems at the dome stage, showed that their shape was often abnormal, with a perturbed cellular organization as the L1 layer showed anticlinal divisions, and divisions in any orientation were observed in the underlying L2 layer (Fig. 2K). To quantify the morphodynamics of CaAMs, we measured their width and height and calculated meristem aspect ratio (height divided by width). Interestingly, we observed on small *dpa4-2 sod7-2* CaAMs (width < 90µm), a higher meristem on average and a more important variability of its shape, compared to WT (Fig. 2M). Larger meristems tended to regain a normal shape when their size increased. We noticed an asynchronous development of the CaAM in *dpa4-2 sod7-2,* in contrast to what was observed in the WT, the size of the meristem was not correlated with the time spent by the plant under LD (Fig. 2N). Nevertheless, both mutant and wild-type meristems switched from the dome to the leaf primordium stage at a similar size. (Fig. 2O). RoAMs showed a delay of initiation between wild-type and *dpa4-2 sod7-2* but no modification of growth dynamics as in CaAMs (Fig. S2E,F). In conclusion, in the *dpa4-2 sod7-2* double mutant CaAM formation is delayed, asynchronous, and associated with an abnormal cellular organisation and shape at the dome stage that reverts to a normal structure at the stage when leaf primordia are initiated.

### The *SOD7*/*NGAL2* and *DPA4*/*NGAL3* genes are required for proper *CUC2* and *CUC3* expression in CaAMs

Because the *NGAL* genes are known negative regulators of the *CUC* gene expression (Engelhorn et al., 2012; Shao et al., 2020), we analysed *CUC2* and *CUC3* expression during CaAM development in *dpa4-2 sod7-2* and WT. Quantitative RT-qPCR showed that *CUC2* and *CUC3* mRNAs levels are increased in developing axillary branches (Fig. S3H-I). To follow *CUC2* and *CUC3* expression during early stages of CaAMs, we introduced the pCUC2:erRFP and pCUC3:erCFP transcriptional reporters into the *dpa4-2 sod7-2* double mutant. In the WT dome stage, pCUC2:erRFP and pCUC3:erCFP reporter expressions were excluded from the meristem and were localized to its base (Fig. 3A,B). In contrast, strong and uniform expression of the reporters was observed in *dpa4-2 sod7-2* domes (Fig. 3C,D). At the leaf primordium stage, pCUC2:erRFP and pCUC3:erCFP reporters were expressed at the boundary domain of the developing leaf primordia in the WT (Fig. 3E,F). A similar expression pattern was observed in the *dpa4-2 sod7-2* mutant, with sometimes weak ectopic expression in the meristem (Fig. 3G,H). Whole mount *in situ* hybridization confirmed a similar localization of *CUC2* and *CUC3* mRNA in the organ primordia boundary domain of both wild type and mutant meristems at the “leaf primordium” stage (Fig. 3M-R). *CUC3* mRNA was distributed throughout the meristem at the *dpa4-2 sod7-2* dome stage, in agreement with the expression pattern of the pCUC3:erCFP reporter (Fig. 3L). *CUC2* mRNA was observed in the rib zone of *dpa4-2 sod7-2* dome stage meristem (Fig. 3K), contrasting with the larger expression of the pCUC2:erRFP reporter (Fig. 3C). Such a reduction of the pattern of *CUC2* mRNA may be due to the post transcriptional regulation of *CUC2* by miR164 (Nikovics et al., 2006; Peaucelle et al., 2007; Sieber et al., 2007). The hypothesis that indeed miR164 may negatively regulated *CUC2* during AM development is supported by the observation that the delay in CaAM development in the *dpa4-2 sod7-2* (t_50_ = 5.74 days) mutant is enhanced by the inactivation of *MIR164A* (t_50_ = 6.77 days for *dpa4-2 sod7-2 mir164a-4*), one of the 3 *MIR164* genes (Nikovics et al., 2006)(Fig. S3A-E). Moreover combining *dpa4-2 sod7-2* with the miRNA resistant version of *CUC2,* CUC2g-m4 (Nikovics et al., 2006) lead to an even stronger phenotype than *dpa4-2 sod7-2 mir164a-4* with no development of CaAM (Fig. S3F-G). Together, these data show that *DPA4* and *SOD7* repress *CUC2* and *CUC3* expression from the developing AM at the dome stage.

**Figure 3.**
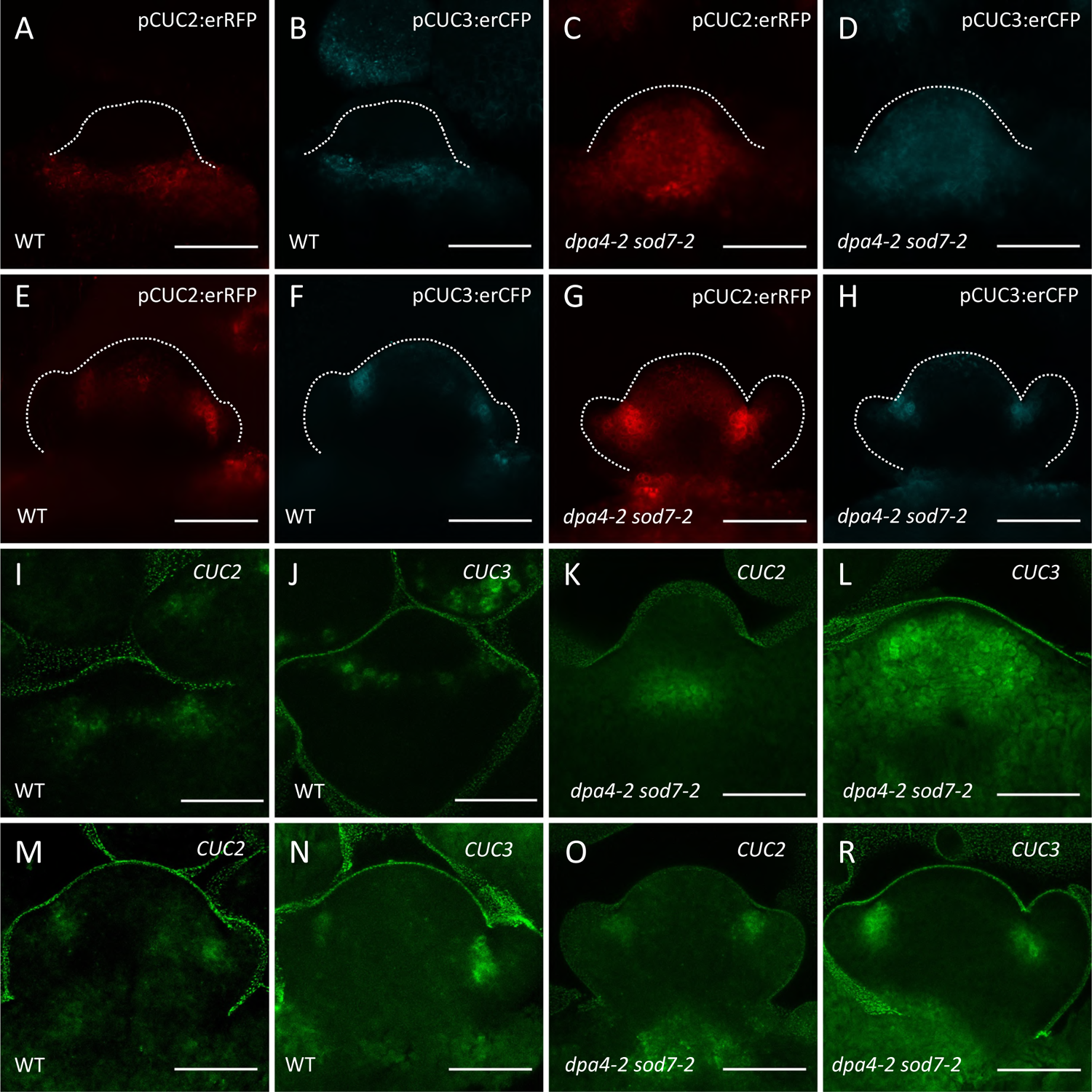
*SOD7* and *DPA4* are required for proper *CUC2* and *CUC3* expression in CaAM (A-H) Maximum projections of tangential optical sections of a pCUC2:erRFP and pCUC3:erCFP reporters in WT and *dpa4-2 sod7-2* during CaAM development at dome stage (A-D) and leaf primordia stage (E-H). (I-R) Maximum projections of tangential optical sections of whole mount *in situ hybridization* of *CUC2* and *CUC3* transcript in WT and *dpa4-2 sod7-2* during CaAM development at dome stage (I-L) and leaf primordia stage (M-R). Plants were grown for 4 weeks in SD conditions and then shifted to LD. Scale bars: (A-R) = 50 µm. The dotted line corresponds to the outline of the meristems and leaf primordia.

### *CUC2* and *CUC3* are required for the delayed CaAM development in the *dpa4-2 sod7-2* double mutant

Because ectopic expression of *CUC2* and *CUC3* coincides with the developmental defects of the *dpa4-2 sod7-2* CaAMs, we next genetically tested the requirement of the *CUC* genes to delay CaAM development in *dpa4-2 sod7-2* (Fig. 4). Introducing the *cuc2-1* (t_50_ = 1.54 days) or *cuc3-105* null allele (t_50_ = 1.52 days) into *dpa4-2 sod7-2* restored growth of the CaAMs (Fig. 4A-J, L, N). The *cuc2-3* weak allele (t_50_ = 1.79 days) also led to a restoration of CaAM development, though to a slightly lower level than the *cuc2-1* null allele (Fig. 4M). In contrast, introducing the *cuc1-13* null allele (t_50_ = 5.57 days) had no effect on CaAM development (Fig. 4G,K). Observation of early stages of CaAM development showed that an active meristem with a proper cellular organization is more rapidly initiated in the *dpa4-2 sod7-2 cuc3-105* triple mutant compared to *dpa4-2 sod7-2* (Fig. 2J lower plot and Fig. 2K, L). Accelerated meristem development has been reported in mutants affected in the strigolactone pathway or the growth repressor *BRC1* (Aguilar-Martínez et al., 2007; Booker et al., 2004; Stirnberg et al., 2002). However, introducing a mutant allele of *BRC1*, *MAX2* or *MAX3* into the *dpa4-2 sod7-2* led to no or weak restoration of CaAM growth (Fig. S4), suggesting that *DPA4* and *SOD7* do not control the strigolactone or BRC1 pathway. Together, these observations suggest that ectopic expression of the *CUC2* and *CUC3* genes is responsible for defective CaAM organization and delayed activity in *dpa4-2 sod7-2*.

**Figure 4.**
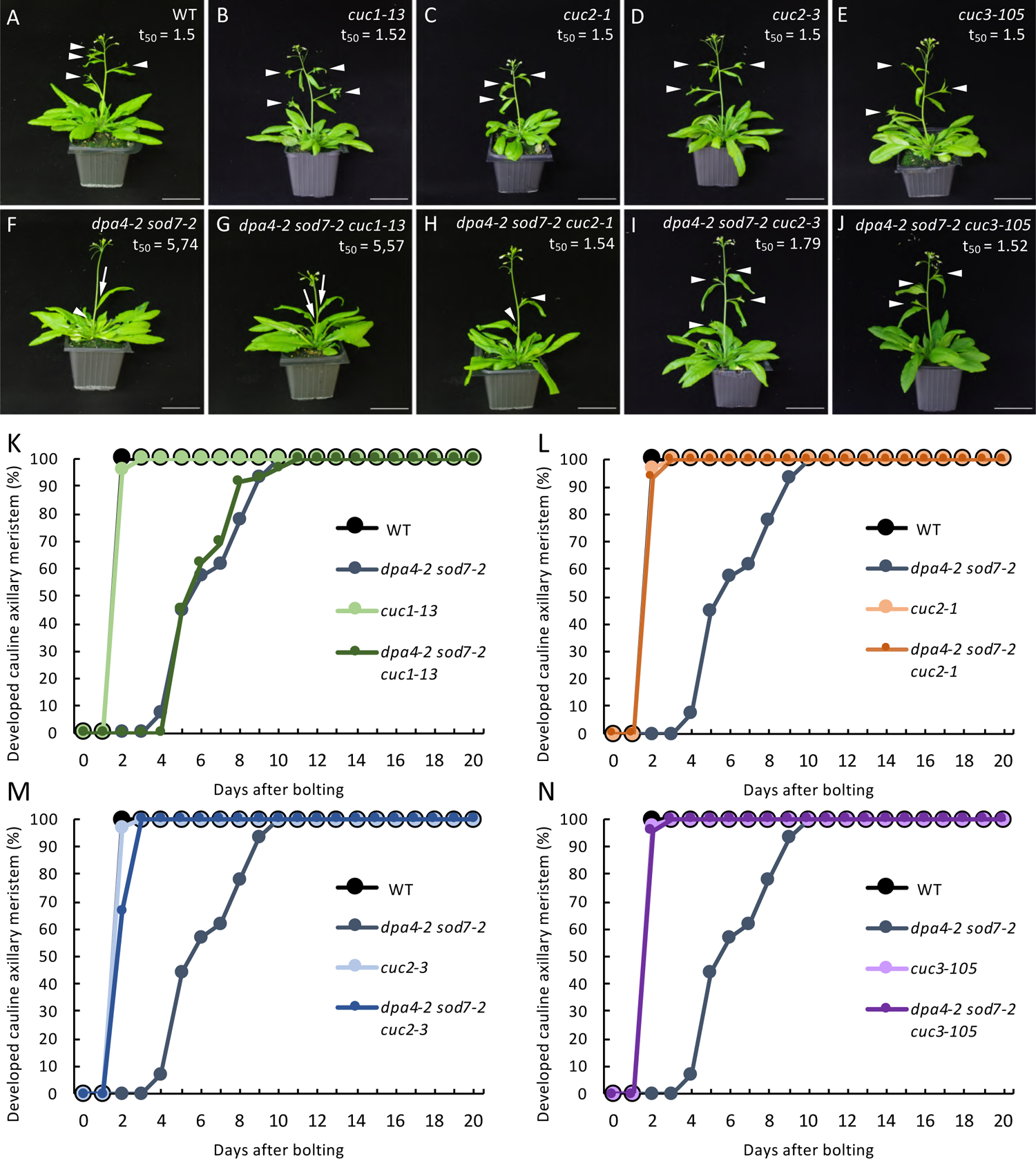
*CUC2* and *CUC3* are required for delayed CaAM development in *dpa4-2 sod7-2* mutants (A-J) Inflorescence of WT, simple *cuc* mutants, double mutant *dpa4-2 sod7-2* and triple mutant *dpa4-2 sod7-2-cuc*. Plants were grown for 5 weeks in LD. White arrowheads point to the developed CaAMs while the arrows point to delayed CaAMs. The time point after bolting at which half of the CaAM are developed (t_50_, in days) is indicated under the genotype. (K-N) Kinetics of CaAM development of WT (K-N), *dpa4-2 sod7-2* (K-N)*, cuc1-13* and *dpa4-2 sod7-2 cuc1-13* (K)*, cuc1* and *dpa4-2 sod7-2 cuc1* (L), *cuc3* and *dpa4-2 sod7-2 cuc3* (M) and *cuc3-105 and dpa4-2 sod7-2 cuc3-105* (N) plants after bolting. Development of the CaAM is indicated as the percentage of developed branches (≥ 3mm) reported to the total number of cauline leaves (*n*≥7). All data were generated in the same experiments, therefore the same WT and *dpa4-2 sod7-2* data were used in panels K to N Scale bars: (A-J) = 5 cm

### *DPA4* and *SOD7* are expressed in the boundary domain and transiently in the stemistem

To follow the expression of the *DPA4* and *SOD7* genes, we generated transcriptional reporters and combined them with the pCUC3:erCFP or pCUC2:erRFP reporters (Fig. 5 and Fig. S5). During early stages, pSOD7:GFP and pDPA4:GFP expression overlapped with pCUC3:erCFP and pCUC2:erRFP in an elongated domain between the meristem and the cauline leaf primordium (Fig. 5A,F,K,P). At the “eye” and “dome” stage, pSOD7:GFP and pDPA4:GFP were maintained in the central domain from which the meristem emerged, while pCUC3:erCFP or pCUC2:erRFP disappeared (Fig. 5B,G,L,Q). At these stages, pDPA4:GFP tended to show a higher expression on the SAM side. Fluorescence quantification along a radial axis from the SAM to the leaf primordium confirmed a stronger depletion of pCUC2:erRFP than pSOD7:GFP in the meristematic dome (Fig. 5C,E,H,J) while pDPA4:GFP showed a peak of expression in the boundary domains closer to the SAM, with a weaker expression in the emerging meristem and on the leaf primordium side (Fig. 5M,O,R,T). Later, pSOD7:GFP and pDPA4:GFP also became excluded from the meristem and limited to the boundary domain where pCUC3:erCFP is expressed (Fig. 5D,I,N,S). A similar dynamic was observed when we compared pCUC2:erRFP with pDPA4:GFP or pCUC3:erRFP with pSOD7:GFP reporters (Fig. S5).

**Figure 5.**
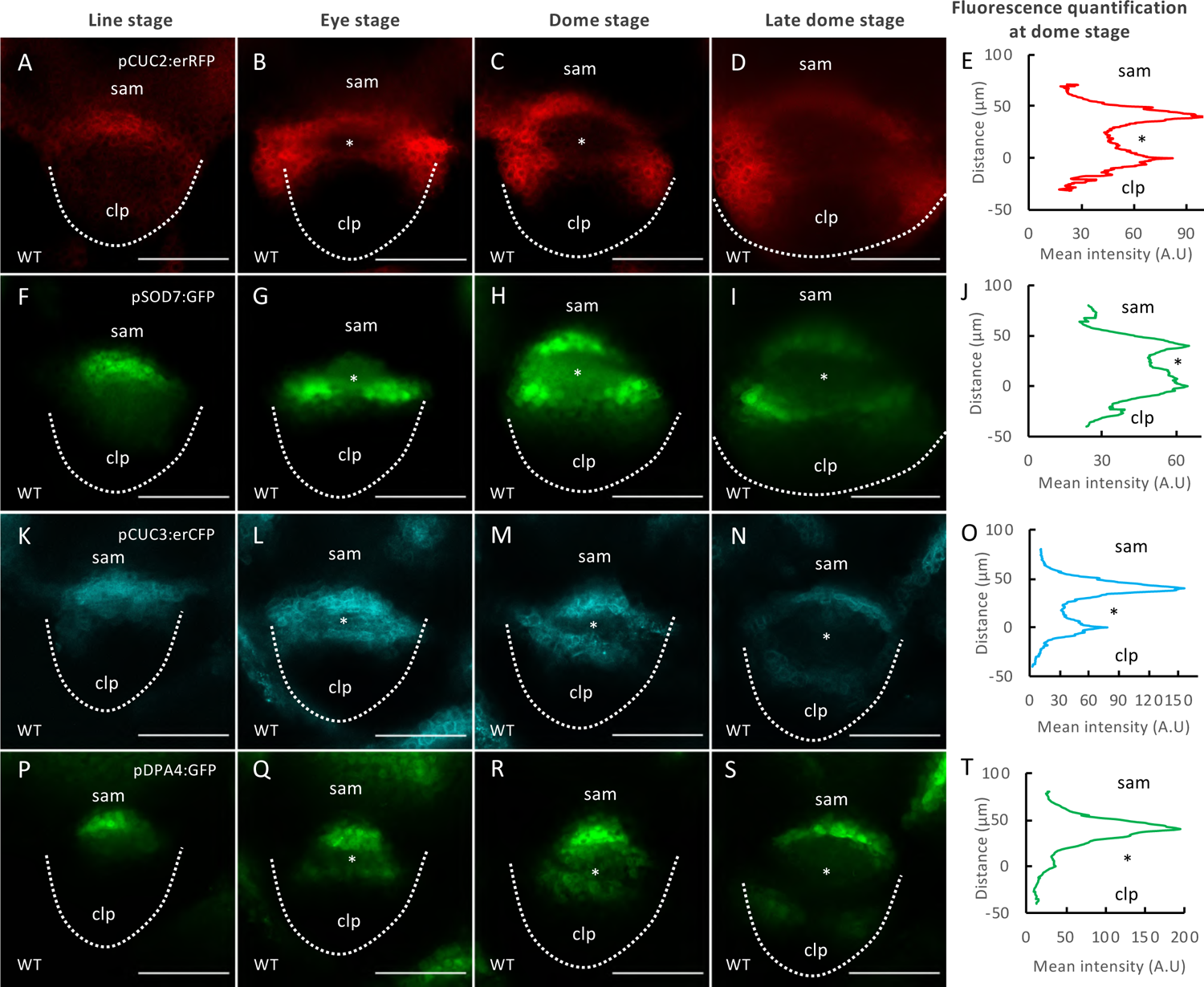
*DPA4* and *SOD7* have overlapping expression patterns with *CUC2* and *CUC3* in the boundary domain and are transiently expressed in the early AM (A-J) Maximum projections of transverse optical sections of plants co-expressing pCUC2:erRFP and pSOD7:GFP reporter lines. Mean fluorescence along the radial axis of CaAM at the dome stage of the pCUC2:erRFP (E) or pSOD7:GFP (J) reporters. (n=6) (K-T) Maximum projections of transverse optical sections of plants co-expressing pCUC3:erCFP and pDPA4:GFP reporter line. Mean fluorescence along the radial axis of CaAM at the dome stage of the pCUC3:erCFP (O) or pDPA4:GFP (T) reporters. (n=6) CaAMs are at the (A,F,K,P) line, (B,G,L,Q) eye,(C,H,M,R) dome and late dome stage (D,I,N,S) Scale bars: (A-P) = 50 µm; sam: shoot apical meristem; clp: cauline leaf primordium; *: AM; The dotted line corresponds to the outline of the cauline leaf primordium.

### Disruption of putative NGAL binding sites in *CUC3* is sufficient to phenocopy the delay of *dpa4-2 sod7-2* secondary stem growth

Next, we investigated the molecular interaction between NGAL proteins and the *CUC* genes. We and others have shown that DPA4 and SOD7 repress *CUC2* and *CUC3* expression and the ABS2/NGAL1 protein directly binds to the *CUC2* promoter (Engelhorn et al., 2012; Shao et al., 2020). It is also known that SOD7 binds to the promoter of the *KLUH* gene through a CACTTG motif (Zhang et al., 2015). RAV1, a transcription factor of the same family as DPA4/SOD7 recognizes a CACCTG motif (Yamasaki et al., 2004) and we found that SOD7 was able to bind *in vitro* to such a sequence present in the *CUC3* promoter (Fig. S6B). Altogether we identified 3 CACTTG and 3 CACCTG motives in the *CUC3* promoter and one CACCTG in the 5’ part of the *CUC3* CDS that could be putative DPA4/SOD7 binding sites (Fig. S6A). In order to test the role of these motifs in *CUC3* expression regulation, we generated a mutated version of *CUC3* with all 7 putative binding sites mutated (pCUC3-6m:CUC3-1m, the mutation in the CDS was silent). We introduced pCUC3-6m:CUC3-1m or a pCUC3:CUC3 control construct in the *cuc3-105* null mutant background. In contrast to what is observed under LD conditions (Fig. 4), *cuc3-105* plants shifted from SD to LD conditions showed a strong defect in CaAM initiation (63% CaAM not initiated at 32LD, Fig. 6A,B). This CaAM initiation defect was suppressed in pCUC3:CUC3 *cuc3-105* (4% CaAM not initiated) and pCUC3-6m:CUC3-1m *cuc3-105* lines (all CaAM initiated) (Fig 6D,E), suggesting that a functional CUC3 was produced from both constructs. However while growth of the secondary stems was similar to the wild type in the complemented pCUC3:CUC3 *cuc3-105* lines, *cuc3-105* lines complemented with the mutated pCUC3-6m:CUC3-1m constructs showed a delayed development of secondary stems similar to *dpa4-2 sod7-2* (Fig. 6A-F). Furthermore, we observed a massive increase of *CUC3* transcript levels in pCUC3-6m:CUC3-m1 *cuc3-105* lines compared to WT, *cuc3-105* and the mutant complemented with pCUC3:CUC3 (Fig. 6G). Those results suggest the putative NGAL binding sites are required to repress *CUC3* expression and *CUC3* overexpression resulting from their mutation lead to a delay in CaAM growth, thus partially phenocopying the *dpa4-2 sod7-2* double mutant.

**Figure 6.**
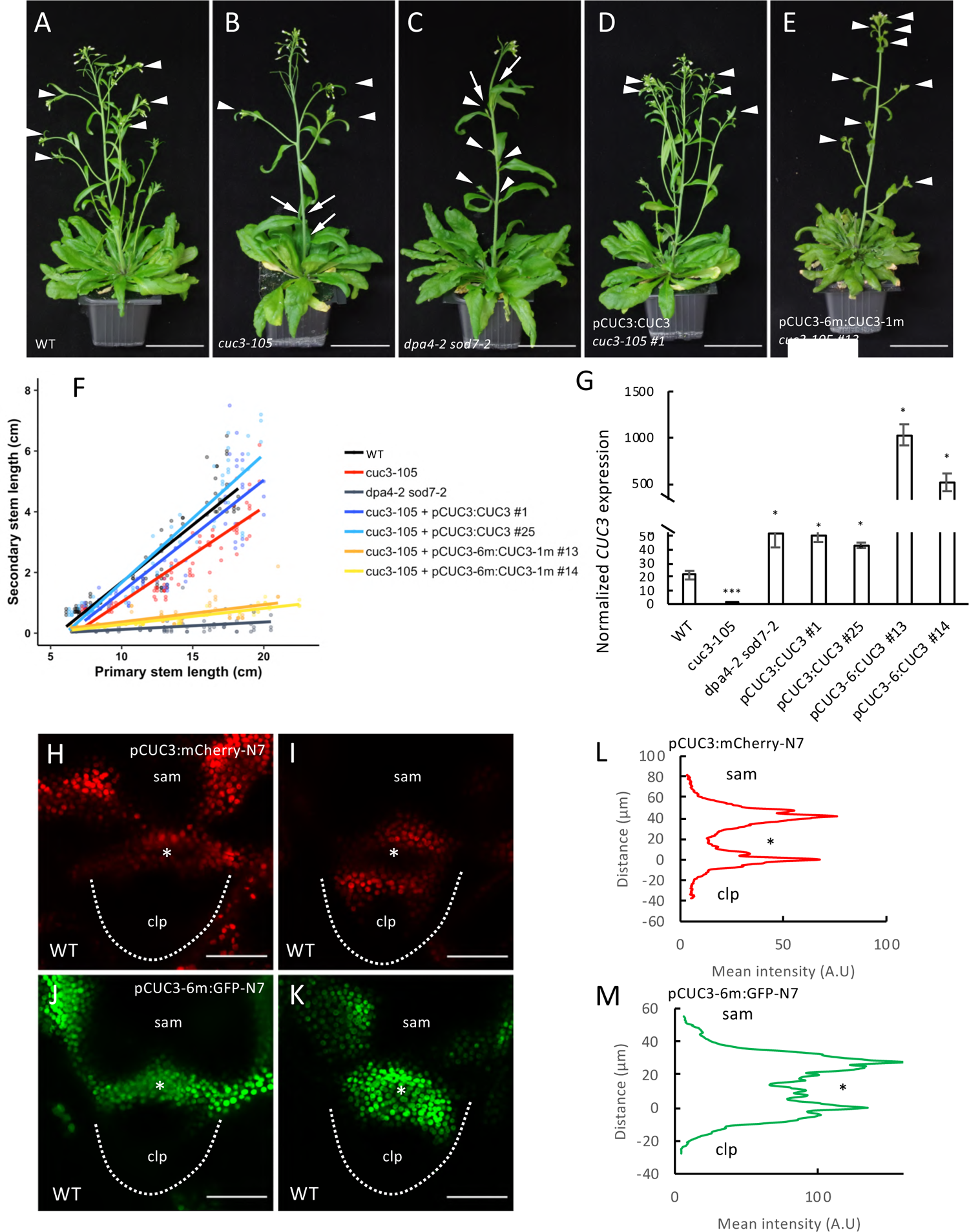
Disruption of putative NGAL binding sites in *CUC3* induces ectopic *CUC3* expression and delay in CaAM development. (A-E) Inflorescence of WT, *cuc3-105* mutant, *dpa4-2 sod7-2* double mutant, pCUC3:CUC3 *cuc3-105* #1 and pCUC3-6m:CUC3-1m *cuc3-105* #13. Plants were grown for 4 weeks in SD and then shifted to LD for 3 weeks. White arrowheads point to the developed CaAMs while the arrows point to delayed CaAMs. (F) Secondary stem length as a function of primary length stem for WT, *cuc3-105* mutant, *dpa4-2 sod7-2* double mutant, pCUC3:CUC3 *cuc3-105* #1 and #25 and pCUC3-6m:CUC3-1m *cuc3-105* #13 and #14. (G) Quantification of the transcript level of *CUC3* by RT-qPCR on 10 day-old seedlings of WT, *cuc3-105* mutant, *dpa4-2 sod7-2* double mutant, pCUC3:CUC3 *cuc3-105* #1 and #25 and pCUC3-6m:CUC3-1m *cuc3-105* #13 and #14. Expressions were normalized using the QREF and REFA genes. A Student’s test was performed to compare the expression levels of mutants in comparison to the wild type (p <0.05 *; p <0.01 **; p <0.001 ***). (H-I) Maximum projections of transverse optical sections of pCUC3:mCherry-N7 or pCUC3-6m:GFP-N7 reporters in wild-type plants during CaAM formation at eye (H,I) and (J,K) dome stage. (L-M) Mean fluorescence along the radial axis of CaAM at the dome stage of the pCUC3:mCherry-N7 (L) or pCUC3-6m:GFP-N7 (M) reporters. (n ≥ 5) Scale bars: (A-E) = 5 cm; (H,I) = 50 µm. sam: shoot apical meristem; clp: cauline leaf primordium; *: AM; the dotted line corresponds to the edge of the cauline leaf primordium.

Because in *dpa4-2 sod7-2* we observed stronger *CUC3* expression than in WT, we next generated a pCUC3-6m reporter line to follow the pattern of the mutated promoter during CaAM development. The control reporter pCUC3:mCherry-N7 showed a clear depletion of the fluorescence in the initiating meristem at the eye and dome stages (Fig. 6H,I), as previously described with the pCUC3:erCFP reporter (Fig. 1). In contrast, the fluorescence of the pCUC3-m6:GFP-N7 reporter remained homogeneous and no clear depletion was observed at eye stage (Fig. 6J) while ectopic fluorescence remained in the developing meristem at the dome stage (Fig. 6K). Quantifications confirmed the diminution of the mean fluorescence intensity inside the dome of in the pCUC3:mCherry-N7 line whereas it remained high in the pCUC3-m6:GFP-N7 line (Fig. 6L-M). At the leaf primordium stage, both wild-type and mutated reporter constructs showed a similar expression in the boundary domain (Fig. S6D,E). This suggests that mutation of putative NGAL binding sites in pCUC3 delays its dynamic repression in the developing meristem. Remarkably, the pCUC3-m6:GFP-N7 reporter has the same dynamic as observed for *CUC3* transcript or pCUC3:erCFP reporter in *dpa4-2 sod7-2.* All these results suggest that disruption of putative NGAL binding sites on *CUC3* can induce ectopic expression of *CUC3* in the center of the CaAM as observed in *dpa4-2 sod7-2,* which in turn delays secondary branch development.

### Repression of the boundary identity is required for stem cell and stem cell niche establishment

Because AM function is associated with *de novo* establishment of stem cells, we next investigated whether stem cell formation is perturbed in *dpa4-2 sod7-2* CaAMs. For this, we first followed the dynamics of a pCLV3:GUS reporter activation in CaAM (Fig. 7A-D). While at 8LD, pCLV3:GUS was expressed in all the wild-type cauline leaf axils, none of the *dpa4-2 sod7-2* double mutant had a visible GUS staining, and at 12LD, only about half of the axils of the double mutant expressed the pCLV3:GUS reporter. The *dpa4-2 sod7-2 cuc3* triple mutant showed a faster pCLV3:GUS activation, confirming that CaAM formation was partly restored in this background compared to the *dpa4-2 sod7-2* double mutant (Fig. 7C,D). To further test whether the delayed pCLV3:GUS was due to the delayed outgrowth of the CaAMs in the double mutant, we compared *CLV3* expression by whole mount *in situ* hybridization in CaAMs of different genotypes at similar morphological stages (Fig. S7A-D). This showed that while at the dome stage most of the wild-type CaAMs expressed *CLV3*, only 23 % of the *dpa4-2 sod7-2* double mutant showed *CLV3* expression (n=17). *CLV3* was restored in all of the dome stage *dpa4-2 sod7-2 cuc3* CaAMs (n=10). This suggested that ectopic expression of *CUC3* in the *dpa4-2 sod7-2* meristem at the dome stage prevents activation of *CLV3*, and that boundary fate needs to be repressed to allow stem cell establishment. Interestingly, when *CLV3* was again observed at the dome stage CaAMs of *dpa4-2 sod7-2* and *dpa4-2 sod7-2 cuc3-105,* its expression pattern was sometimes abnormal as *CLV3* tended to be expressed in the centre of the meristem as was observed during wild-type RoAM initiation (Fig. S7E,F, Fig. 1S). Respectively 78% and 70% of *dpa4-2 sod7-2* (n=14) and *dpa4-2 sod7-2 cuc3-105* (n=10) CaAMs showed such central ectopic expression of *CLV3*. This ectopic central expression of CLV3 is likely to be a transition phase as it was mostly observed on small CaAM in *dpa4-2 sod7-2* (width <90µm), while larger meristems showed a normal apical expression pattern (Fig. 7E). Interestingly, the ectopic expression of *CLV3* in *dpa4-2 sod7-2* can be correlated with the perturbed cellular organization observed at the dome stage in *dpa4-2 sod7-2* (Fig. 2).

**Figure 7.**
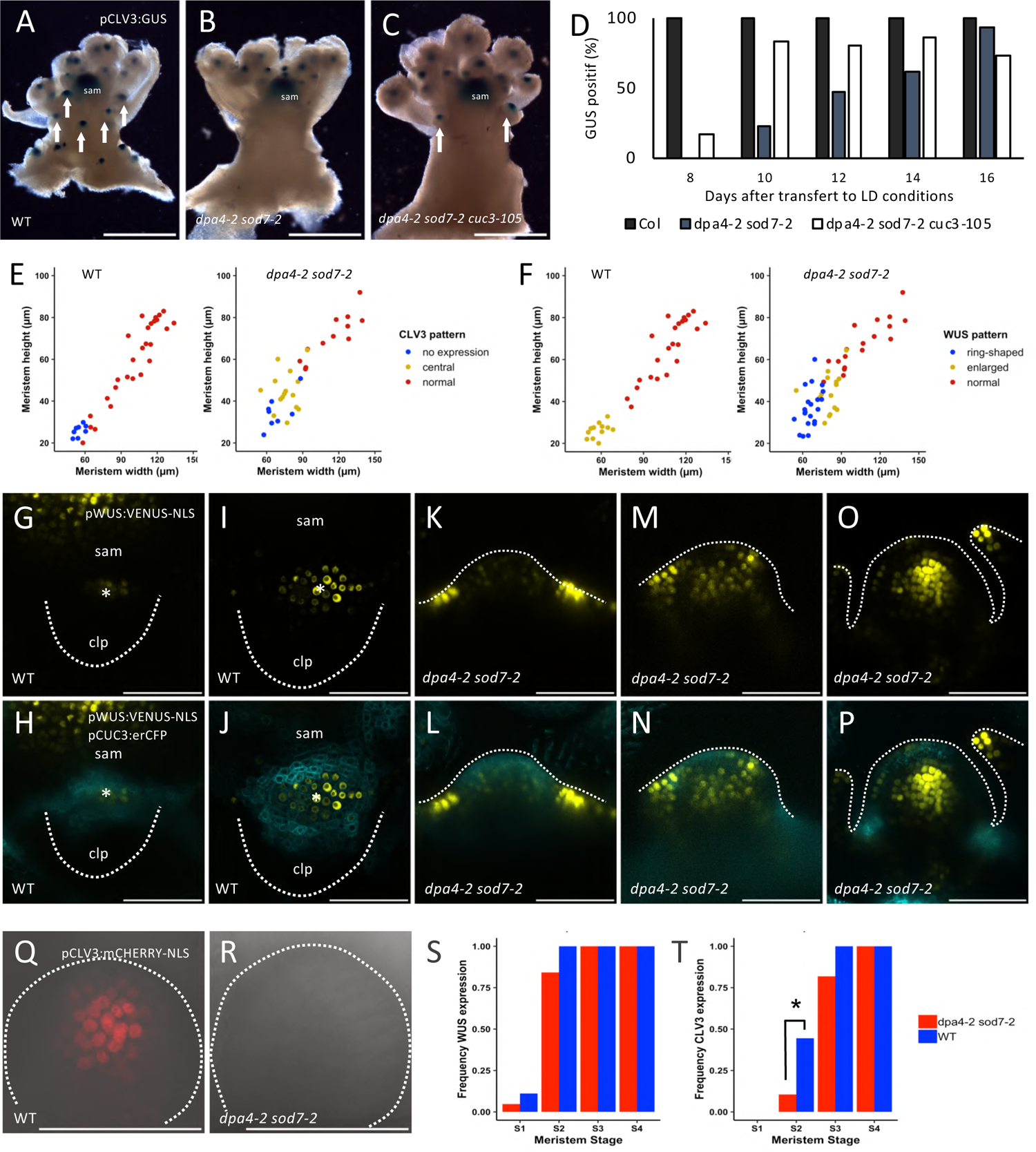
Stem cell specification is delayed in *dpa4-2 sod7-2* AM and floral meristems. (A-C) Expression of a pCLV3:GUS reporter in (A) *dpa4-2 sod7-2* (B) *dpa4-2 sod7-2 cuc3-105* (C) inflorescences. Plants were grown for 4 weeks in SD and then shifted to LD for 10 days. The arrows point CaAM with pCLV3:GUS expression and sam indicate the shoot apical meristem (D) Quantification of GUS positive CaAM with pCLV3:GUS expression in plants shifted to LD. (E) *CLV3* expression pattern as a function of CaAM width and height in WT and *dpa4-2 sod7-2*. “normal” is CLV3 expressed in the apical region as shown in Fig S7A,C,D,E, while “central” is CLV3 expression in the centre of the meristem as shown in Fig S7F. (F) *WUS* expression pattern as a function of CaAM width and height in WT and *dpa4-2 sod7-2*. “enlarged” is *WUS* expressed in the entire meristem as shown in panels J and N, “ring-shaped” is *WUS* expressed at the base of the meristem as in panel L, and “normal” is WUS expressed in few cells in the centre of the meristem as in panel P. (G-P) Maximum projections of transverse (G-J) and tangential (K-P) optical sections of pWUS:VENUS-NLS and pCUC3:erCFP in wild-type and *dpa4-2 sod7-2* during CaAM development. Plants were grown for 4 weeks in SD conditions and then shifted to LD. (Q-P). Maximum projections of transverse optical sections pCLV3:mCHERRY-NLS in floral meristems at stage 2 in WT and *dpa4-2 sod7-2*. (S,T) *WUS* and *CLV3* expression as a function of floral meristem stage. A Fisher’s test was performed to compare the expression levels of the mutants in comparison to the wild-type (p <0.05 *). Scale bars: (A-C) = 0.5cm; (G-R) = 50µm; sam: shoot apical meristem; clp: cauline leaf primordium; *: AM; the dotted line corresponds to the outline of the cauline leaf primordium (G-J), AM (K-P) or floral meristem (Q,R)

Next, because *WUS* is expressed earlier and activates *CLV3*, we wanted to know if *WUS* expression was also delayed in *dpa4-2 sod7-2.* For this, we compared the dynamics of the pWUS:VENUS-NLS and pCUC3:erCFP reporters in wild-type and *dpa4-2 sod7-2* plants (Fig. 7G-P). In the wild-type background, at the eye-stage, we observed a few cells expressing pWUS:VENUS-NLS in the center of the developing CaAM where pCUC3:erCFP expression started to disappear (Fig. 7G,H). Later on during the dome stage, pWUS:VENUS-NLS expression pattern enlarged and was highest in the meristem part where pCUC3:erCFP expression was low (Fig. 7I,J). Interestingly, in the smallest *dpa4-2 sod7-2,* CaAM pWUS:VENUS-NLS was very strong in a few cells at the outer base of the meristem, forming a ring-shaped structure which was complementary to the pattern of pCUC3:erCFP inside the whole dome of the CaAM (Fig. 7K,L). Much weaker pWUS:VENUS-NLS expression was detected in a few cells within the meristem. Later on, pWUS:VENUS-NLS expression increased in the meristem of *dpa4-2 sod7-2* mutants (Fig. 7M,N). Lastly, during leaf primordium stage, a normal expression of pWUS:VENUS-NLS was observed in *dpa4-2 sod7-2* while pCUC3:erCFP expression returned to the boundary domains (Fig. 7O,P). Whole mount *in situ* hybridization confirmed ectopic *WUS* expression at the base of the *dpa4-2 sod7-2* meristems while the *dpa4-2 sod7-2 cuc3-105* triple mutant showed a wild-type *WUS* pattern (Fig. S7G-I).

Linking these *WUS* patterns with meristem size, confirmed that in the wild type, small meristem had an enlarged *WUS* expression while at later stages it became restricted to the centre of the meristem (Fig. 7F). In *dpa4-2 sod7-2* mutants, *WUS* switched from an initial expression in ring-shaped pattern around its base to an expression throughout the meristem before becoming restricted to a central normal domain (Fig. 7F). Those results suggested that ectopic expression of *CUC2/CUC3* prevents activation of *WUS* in the meristem.

Together, our results lead to a scenario where the DPA4 and SOD7 transcription factors are essential for a rapid repression of the *CUC2*/*CUC3* genes from the developing AM during the expansion phase in which the number of meristematic cells increases. If such a rapid repression does not occur, ectopic CUC2/CUC3 expression would lead to defective meristem growth and organisation, and delayed activation of *WUS* in the meristem, which in turn would lead to a delayed activation of *CLV3* and hence to defective *de novo* stem cell niche establishment.

### *DPA4 and SOD7* facilitate the establishment of the stem cells in the floral meristem

To test whether *DPA4* and *SOD7* had a general role in *de novo* stem cell formation we analysed stem cell establishment in newly formed floral meristems using the pWUS:VENUS-NLS and pCLV3:mCHERRY-NLS reporters. In agreement with previous reports (Mayer et al., 1998), in the wild type, pWUS:VENUS-NLS was expressed in a small proportion of the floral meristems at stage 1 and was expressed in all stage 2 flowers (Fig. 7S). Slightly less stage 1 and stage 2 *dpa4-2 sod7-2* floral meristems expressed pWUS:VENUS-NLS, suggesting a small delay in *WUS* activation which was also observed when the meristems were staged according to their size (Fig. 7S). Interestingly, *CLV3* expression was more affected than *WUS.* Indeed, while 44% of wild-type stage 2 floral meristems expressed pCLV3:mCHERRY-NLS, only 11% of the *dpa4-2 sod7-2* expressed it (Fig 7Q,R,T). At stage 3, all wild-type meristems expressed the CLV3 reporter while it was absent from 18% of the *dpa4-2 sod7-2* meristems (Fig. 7T). Accordingly, pCLV3:mCHERRY-NLS started to be expressed in *dpa4-2 sod7-2* floral meristems that were almost twice as big as the wild type (Fig. S8). Based on those results, we can conclude that *DPA4* and *SOD7* act together to facilitate *de novo* stem cell establishment in floral meristems.

## DISCUSSION

Stem cells are important throughout the life of all living organisms and, in plants, new population of stem cells and their enclosing meristems have to be formed throughout their life to enable continuous growth and branching. Such meristems are formed in the axils of leaves from boundary domains that maintain meristematic features. Work in the recent years has shown that AM initiation requires the maintenance of a meristematic fate by a dense network of interacting transcription factors and hormones, in which the *CUC* boundary genes play a central role, and accordingly *cuc* mutants show strong defects in meristem initiation (Hibara et al., 2006; Keller et al., 2006; Müller et al., 2006; Raman et al., 2008; Tian et al., 2014). Here we show that the expression of the *CUC* genes has to be down-regulated for the initiating meristem to proceed to the establishment phase and become active. We show that the NGAL transcription factors DPA4 and SOD7 are required to effectively remove *CUC* expression from the initiating AM. *CUC* mis-expression in the developing AM leads to asynchronous and delayed meristem formation, associated with abnormal cellular organization. Notably, ectopic expression of these boundary cell fate genes prevents stem cell establishment that is required for meristem activity. Because we observed that delayed stem cell formation also occurs in floral meristems of the *dpa4-2 sod7-2* double mutant, our work reveals a conserved genetic circuit by which the NGAL transcription factors repress the *CUC* boundary genes to allow *de novo* stem cell establishment in newly formed meristems.

Arabidopsis can form AMs from both its rosette and cauline leaves and our work highlights differences previously unknown between the development of these two structures. First, while the formation of the RoAMs is a slow process extending over numerous plastochrons, the formation of the CaAM is much faster. For instance, *WUS* expression is initiated in P13 in RoAMs (Wang et al., 2017) while we observed *WUS* expression as early as P7 in CaAMs. As a consequence, the balance between relative growth of the leaf and the associated AM is pushed towards the leaf in the rosette and towards the meristem in cauline leaves. Indeed, we observed within successive CaAMs a trend of the AM to develop even faster relative to the leaf primordium in the upper nodes before the reproductive stage. Interestingly, it has been suggested that in the case of the floral meristem (a modified AM), the growth of a cryptic bract (a modified leaf) is suppressed (Long and Barton, 2000; Ohno et al., 2004). Altogether, this suggests that bract suppression during flower development may not be such an abrupt event as previously thought but could be the culminating point of a progressive reduction of lateral organ growth relative to AM development as the plant further matures.

A second difference between RoAMs and CaAMs, is that CaAMs grow out directly after their initiation with no apparent phase of dormancy. As a consequence, mutations in genes inhibiting AM outgrowth such *BRC1* or those of the strigolactone pathway like *MAX2*/3 (Aguilar-Martínez et al., 2007; Booker et al., 2004; Stirnberg et al., 2002) do not further increase CaAM branching. Our genetic analysis indicate that the slow outgrowth of the *dpa4-2 sod7-2* CaAMs can be slightly sped-up by mutations in the strigolactone pathway components or *brc1*, suggesting that these pathways may still be active in CaAMs. However, the level of phenotypic restoration observed in these mutants is much lower than the one observed with the *cuc* mutations, suggesting that these pathways are not the ones primarily affected in the *dpa4-2 sod7-2* mutants.

A third difference between CaAMs and RoAMs can be seen in the dynamics of gene activation leading to stem cell establishment. While in both organs, *WUS* is activated before *CLV3*, in CaAM, the *WUS* domain shifts from an apical to a central position while in RoAM, *WUS* is already expressed in the central domain. In turn, *CLV3* is properly positioned in apical position from the beginning in CaAM, while in RoAM it moves from a central to an apical position. (Xin et al., 2017). Further characterizing in CaAM cytokinin signaling or HAM gene spatial patterns, that have been shown to contribute to stem cell establishment in RoAMs, will be necessary to understand these differences (Han et al., 2020a, 2020b; Wang et al., 2017; Zhou et al., 2018).

Our data show that while *CUC* genes are required for AM formation (Hibara et al., 2006; Raman et al., 2008), likely by preventing cell differentiation and maintaining cells in a meristematic fate, their expression has to be negatively regulated to allow proper meristem establishment. Their prolonged, ectopic expression in the meristem is associated with asynchronous growth, abnormal cellular and meristem organization and delayed organ initiation. These defects can be traced back to some roles of the *CUC* genes as these genes have been shown to affect cell proliferation and cell expansion (Kierzkowski et al., 2019; Larue et al., 2009; Peaucelle et al., 2007; Serra and Perrot-Rechenmann, 2020; Sieber et al., 2007) as well as auxin transport and signaling (Bilsborough et al., 2011; Heisler et al., 2005; Maugarny-Calès et al., 2019). However, following an initial phase during which *dpa4-2 sod7-2* meristems are misshapen, they recover, restraining *CUC2* and *CUC3* to the boundary. Because this transition is accelerated in a *cuc3* mutant background, it suggests that ectopic *CUC* activity may be limiting for this. Such a reversion to a recovering meristem could be controlled by genetic factors. For instance, *ABS2*, the third *NGAL* gene, may contribute to exclude *CUC* expression from the meristem. However, because no major differences were observed between AM phenotype in *dpa4-2 sod7-2* double and *dpa4-2 sod7-2 abs1* triple mutant, this suggests *ABS2* role may be limited. Alternatively, miR164, which is a well-known repressor of *CUC2* expression that acts independently of *NGAL* genes (Engelhorn et al., 2012) may also be involved. Our genetic analysis with mutations modifying miR164 activity supports such a role.

An alternative hypothesis also emerges from the comparison with the patterning of the leaf margin that leads to teeth formation. In the case of the leaf margin, a pattern with discontinuous *CUC* expression stripes forms as an emergent property of interconnected feedback loops between *CUC* activity and auxin transport and signalling (Bilsborough et al., 2011). In addition to the dynamics of these feedback loops, growth is essential for this patterning process as it generates a cellular template large enough for the feedback loops to be deployed. In such a view, *CUC* expression patterns would be able to reorganize once the slowly growing meristems of the *dpa4 sod7* mutants would reach a critical size threshold. Testing such an hypothesis would require further investigations of the interconnections between AM growth and gene expression dynamics for instance through combined modelling and experimental perturbation of growth.

The final step in meristem formation is the *de novo* establishment of an active stem cell niche. This is essential for the indeterminate fate of AM but is also required for proper floral morphogenesis as a reduction of the inner organs is observed in *wus* flowers in which the stem cell niche is not properly specified (Laux et al., 1996). In both axillary and floral meristems, *WUS* activation precedes the expression of the stem cell marker *CLV3*. Here, we show that in the wild-type initiating CaAM, *WUS* expression is rapidly induced in a few cells that are depleted for *CUC3* expression. Later, the *WUS* domain progressively enlarges, occupying most of the developing meristem that is complementary to the *CUC3*-expressing cells. In *dpa4-2 sod7-2*, *CUC2* and *CUC3* mis-expression during the dome stage profoundly modifies *WUS* expression patterns, which becomes mostly restricted to a ring-shaped structure at the base of the meristem and excluded from the meristem itself. Therefore, as in the wild type, the expression patterns of the *CUC* genes and *WUS* are essentially mutually exclusive in *dpa4 sod7* double mutant. This observation suggests a scenario in which *CUC3* represses *WUS* expression although alternative scenarios are possible. For instance, it has been suggested that geometrical changes of an emerging meristem may be sufficient for the activation of new *WUS* and *CLV3* domains (Gruel et al., 2016). In such a view, defects in *WUS* and *CLV3* activation in the double *dpa4-2 sod7-2* mutant could be a consequence of abnormal meristem growth or shape.

While AM are initiated from a group of cells expressing the *CUC2* and *CUC3* organ boundary genes, these boundary domains are located on one side of the initiating floral meristem (Heisler et al., 2005). Indeed, in floral meristems, *CUC* genes are expressed at stage 1 forming the boundary between the floral primordia and the SAM until their expression disappears at stage 4 (Hibara et al., 2006). Despite these differences in the origin of the meristem relative to the boundary domain, *dpa4 sod7* mutants show a delayed stem cell specification in both AM and floral meristems, suggesting that the *NGAL*/*CUC* regulatory module similarly controls *de novo* stem cell formation in all aerial post-embryonnically formed meristems.

## MATERIALS & METHODS

### Plant material and growth conditions

All genotypes are in the Columbia-0 (WT) ecotype. *The cuc2-1* mutant was isolated from Landsberg *erecta* ecotype but was backcrossed 5 times in Col-0 (Hasson et al., 2011). The mutant allele, *dpa4-2* (Engelhorn et al., 2012)*, sod7-2* and *dpa4-3* (Zhang et al., 2015), *abs1* (Shao et al., 2012), *cuc1-13, cuc2-3*, *cuc3-105* (Hibara et al., 2006), *brc1-2* (Aguilar-Martínez et al., 2007), *max2-1* (Stirnberg et al., 2007) and *max3-11* (Booker et al., 2004) were previously described, as well as the pCUC3:erCFP (Gonçalves et al., 2015), pCUC2:erRFP (Gonçalves et al., 2017), CUC2g-m4 (Nikovics et al., 2006), pCLV3:GUS (Brand et al., 2002) and pCLV3::mCHERRY-NLS/pWUS::3X VENUS-NLS (Pfeiffer et al., 2016).

Seeds were soaked in water at 4 °C for 48 hours prior to sowing. Plants were grown in soil either in long-day (LD) conditions [2 h dawn (19°C, 65% hygrometry, 80 μmol.m-2.s-1 light), 12h day (21 °C, 65% hygrometry, 120 μmol.m-2.s-1 light), 2h dusk (20 °C, 65% hygrometry, 80 μmol.m-2.s-1 light), 16 h dark (18 °C, 65% hygrometry, no light)] or in short-day (SD) conditions [1 h dawn (19 °C, 65% hygrometry, 80 μmol.m-2.s-1 light), 6 h day (21 °C, 65% hygrometry, 120 μmol.m-2.s-1 light), 1 h dusk (20 °C, 65% hygrometry, 80 μmol.m-2.s-1 light), 16 h dark (18 °C, 65% hygrometry, no light)] and then shifted to LD. Seedlings from Fig. 7G were grown in vitro on Arabidopsis medium Duchefa in long day conditions [16h light / 8h dark at 21 °C].

### Enhanced Yeast One-Hybrid Analysis

CUC2 and CUC3 promoters were amplified by PCR using promCUC2 Fwd and promCUC2 Rv (3.7 kb) and prCuc3 – Fw and prCuc3 – R (4.3 kb) (see Primers in Supplemental Table 1). They were recombined with the 5’TOPO plasmid and then into pMW2 and pMW3 for HIS3 and LACZ reporter selection, respectively. Bait constructs were transformed into yeast as described in Gaudinier et al. (2011) and selected for on -His and -Ura dropout media and for minimal auto-activation in the reporter assays. The prey transcription factor collection used is described in Gaudinier et al. (2011) and Truskina et al. (2021) (see full list in Supplemental Table 2). Bait and prey transcription factors were introduced into a diploid yeast colony using the mating method as described in Gaudinier et al. (2011). The interaction between SOD7 and pCUC3 led to LACZ reporter activation but no HIS3 activation.

### Generation of transgenic plants

2.8 kb promoter of *DPA4* was amplified with Pdpa4-2FW and Pdpa4-2RV (Supplemental Table 1) and inserted in front of a GFP in the pMDC107 to generate pDPA4:GFP. 2.1 kb promoter of *SOD7* (Zhang et al., 2015) was amplified with SOD7Profwattb1 and SOD7Prorvattb2 primers and inserted in front of a GFP in the pMDC107 to generate pSOD7:GFP. The promoters of *DPA4* and *SOD7* were cloned using a Gateway strategy.

All the parts used by a Goldenbraid 2.0 strategy (Sarrion-Perdigones et al., 2013) are listed in Supplemental Table 3. 4.3 kb promoter of *CUC3* was amplified and domesticated with GB_S1pCUC3S2_F and GB_S1pCUC3S2_R and inserted in the pUPD2. 3 patches of *CUC3* coding sequence of respectively 175bp, 644bp and 273bp were amplified with CUC3_S2F and CUC3_dom1R for patch1, CUC3_dom1F and CUC3_dom2R for patch2 and CUC3_dom2F and CUC3_S7R for patch3, combined to obtain a 1kb fragment and then inserted in the pUPD2. To generate *CUC3-1m*, we used *CUC3* pUPD2 as a matrix and amplified with CUC3_S2F and CUC3_CDS_PF3_r a first patch and with CUC3_CDS_PF3_f and CUC3_S7R a second patch to generate a silent mutation into the *NGAL* binding site mutation BS3 (Fig. S7A). A 3.7 kb fragment of *CUC3* promoter with the six binding sites mutated (pCUC3-6m) (Fig. S7A) was synthesized by Genewiz (https://www.genewiz.com/) in a pUC-GW-Kan vector. Then the 3,7 kb pCUC3-6m fragment was excised from pUC-GW-Kan with NsiI-PstI enzymes and inserted into the pCUC3 pUPD2 vector also digested NsiI-PstI enzymes to generate a pCUC3-6m pUPD2.

To form the transcriptional unit (T.U), the different parts into the pUPD2 vectors were inserted in an pDGB3_α1 binary vector. The differents T.U in pDGB3_α1 were combined either with pnos:hygro:tnos pDGB3_α2 or with pCMV:DSRed:tnos pDGB3_α2 into an pDGB3_Ω1 binary vector.

The resulting constructs (pMDC107 or pDGB3_Ω1) were sequence-verified and transferred into Agrobacterium tumefaciens strain GV3101. Plants were transformed by floral dipping. Primary transformants were selected *in vitro* for their resistance to hygromycin (pMDC107, pDGB3_Ω1) or selected with the red selection marker (pDGB3_Ω1). Several primary transformants were analysed for their phenotype and for each construction at least two independent lines were selected based on resistance segregation.

### RNA whole mount *in situ* hybridization

RNA *in situ* hybridization was completed as described in Chelysheva et al., (in preparation). Primers used to amplified the probes are indicated in Supplemental Table 1*. In situ* signal was revealed using the Vector® Blue Substrate Kit, Alkaline Phosphatase (Vector Laboratories) and imaged by confocal microscopy (see Supplemental Table 4)

### CaAM preparation for confocal imaging

Plants were grown for 4 weeks in SD and shifted to LD. All observations were done in CaAM between 5 and 16 days after shifting in LD. All the observations were on fresh samples except for Fig. 1A-D, 2K,L where samples were fixed on 4% paraformaldehyde under vacuum for 1h and clearing in Clearsee (xylitol 10%, urea 25%, deoxycholate15%) (Kurihara et al., 2015) and Calcofluor (0.1%) for at least 2 weeks. Hand dissected meristems were mounted between slide and coverslip with Tris HCl 10mM pH = 8,5, Triton 0,01%.

### Confocal imaging

Confocal imaging was performed on a Leica SP5 inverted microscope (Leica Microsystems, Wetzlar, Germany). Lenses are Leica 40x HCX PL APO CS. Acquisition parameters are presented in Supplemental Table 4. Imaging was done from above for apices until 10-12 LD while older apices had to be imaged from the side. Figures were made using ImageJ and FigureJ (Mutterer and Zinck, 2013). All the confocal images are maximum projections.

### Signal normalisation and averaging

Fluorescence profiles were computed using Fiji, then spatially normalized and averaged based on the two major signal peaks. First, each peak localization was determined along each individual signal profile. For this, the profile was split into two, on either side of the profile median position. Each of the two peaks was localized as the position of the maximal signal value on the corresponding side. To register several profiles, resulting peaks were put in correspondence using linear scaling and translation of the profile axis. In the resulting referent axis, the distance between the two peaks can either be arbitrary chosen, e.g., by specifying that the two normalized peaks are separated one from each other by a distance of 1 unit, or automatically from input data, e.g., by using the average distance between peaks computed from the data. After individual data normalization, profiles were averaged to yield the mean signal intensity profile. A script was developed in R for this and used to generate Fig. 5E,JO,T and Fig. 6 L,M

### Scanning electron microscopy

Freshly sampled tissues were cooled to −33C° by a peltier cooling stage (Deben) and observed with a Hirox SH-1500 benchtop scanning electron microscope.

### RNA extraction and RT-qPCR expression analysis

Total RNA were isolated using RNAeasy Plant Mini Kit (Qiagen) following manufacturer’s instructions for plant tissue including DNAse treatment. Reverse transcription was performed using RevertAid H Minus M-MuLV Reverse transcriptase (Fermentas) followed by a RNAse H treatment for 20 min at 37°C to eliminate DNA-RNA duplexes. Real time PCR analysis was performed on a Bio-Rad CFX connect machine using the SsoAd-vance Universal SYBR Green Supermix following manufacturer’s instruction. PCR conditions are as follows: Conditions: 95 °C 3min; (95 °C 10s; 63 °C 10s; 72 °C 10s) x45 cycles. Primers used for real time PCR analysis are available in Supplemental Table 1. Expression data were normalized using the ΔΔCt method (Livak and Schmittgen, 2001).

### GUS staining

GUS staining was performed as described (Sessions et al., 1999) in the presence of 0.2 mM potassium ferricyanide and potassium ferrocyanure. The reaction was stopped with 95% ethanol, which was also used to remove the chlorophyll from the tissues.

### Phenotypic analysis

A count of CaAM and RoAM development was carried out over a period of twenty days after bolting (determined when the primary stem > 1 mm) on plants grown 5 weeks on LD. CaAM and RoAM were counted every 2 days. A meristem is considered present when it begins to grow and be sufficiently visible to the naked eye (> 3mm). In addition, the final number of stem leaves and rosettes was also counted. We calculated the time point after bolting at which half of the CaAMs or RoAM were developed (t_50_) using a R script.

A count of the stages of development of CaAM was carried out over a period of twenty days after bolting (determined when the primary stem > 1 mm) on plants grown 4 weeks in SD and then shifted to LD. Observations were done on CaAM between 8 and 28 days after shifting in LD using a binocular microscope.

## ACKNOWLEDGMENT

We thank P. Cubas, C. Rameau and the NASC for providing seeds. We thank N. Arnaud, N. Bouré, M. Azzopardi, L. Gissot for providing parts used in the Goldenbraid cloning steps. We thank members of the FTA team at IJPB for discussion and N Arnaud for comments on the manuscript. The IJPB benefits from the support of Saclay Plant Sciences-SPS (ANR-17-EUR-0007). This work has benefited from the support of IJPB’s Plant Observatory technological platforms and financial support from the France Berkeley Fund.

## AUTHOR CONTRIBUTION

AN, PL and AMC conceived the project and PL supervised the project. AN performed most of the experiments with the help of PL. AMC, AMB and MS performed the Y1H screen under the supervision of SB. AMC did the preliminary genetic analysis. BA contributed to the generation of the double mutant and transgenic lines. LC conceived the whole mount *in situ* protocol and supervised AN for this. Yu.L performed the gel shift experiment under the supervision of YL. JB wrote the fluorescence average script. AN and PL wrote the paper with inputs of AMC.

## LEGENDS TO THE SUPPLEMENTAL FIGURES

**Figure 1 Supplemental. *WUS* and *CLV3* expression in CaAM and RoAM** (A-C) Maximum projections of radial optical sections of a pWUS:VENUS-NLS (A) and pCLV3:mCHERRY-NLS (B) reporter lines and the overlay (C) during CaAM formation. (D-F) Maximum projections of radial optical sections of a pWUS:VENUS-NLS (D) and pCLV3:mCHERRY-NLS (E) reporter lines and the overlay (F) during RoAM formation. The number of days in LD conditions is indicated. Scale bars: (A-F) = 50 µm

**Figure 2 Supplemental. *DPA4* and *SOD7* are required for rapid development of cauline AMs.** (A,B) Kinetics of CaAM or RoAM development of all *ngal* simple and multiple mutants after bolting. Development of the meristems is indicated as the percentage of developed branches (≥ 3mm) reported to the total number of cauline or rosette leaves (*n*≥11). (C,D) SEM observations of WT and *dpa4-2 sod7-2* RoAM from leaf 7 on plants grown 4 weeks in SD. (E) Quantification method for (F) on a maximum projection of transverse optical sections of pCUC3:erCFP reporter in WT SAM on plants grown 4 weeks in SD. The red line represents the width of the RoAM and the blue line the distance between the SAM and the RoAM. (F) RoAM width as a function of the distance between the same RoAM and the SAM. (G,H) Inflorescence of WT and *dpa4-3 sod7-2* double mutants. Plants were grown for 5 weeks in LD. White arrowheads point to the developed CaAMs while the arrows point to delayed CaAMs. Scale bars: (C,D) = 200 µm; (E) = 100 µm, (G,H) = 5cm

**Figure 3 Supplemental. Genetic interaction between *MIR164* and *DPA4*/*SOD7* during CaAM development and *CUC2*/*CUC3* mRNA quantification in *dpa4-2 sod7-2*.** (A-D) Inflorescence of WT, and *mir164a-4*, *dpa4-2 sod7-2* and *dpa4-2 sod7-2 mir164a-4* mutants. Plants were grown for 5 weeks in LD. White arrowheads point to the developed CaAMs while the arrows point to delayed CaAMs. The time point after bolting at which half of the CaAM are developed (t_50_) is indicated under the genotype. (E) Kinetics of CaAM development after bolting. Development of the CaAM is indicated as the percentage of developed branches (≥ 3mm) reported to the total number of cauline leaves (*n*≥8). (F) Inflorescence of *dpa4-2 sod7-2 cuc2g-m4* mutant. Plants were grown for 6 weeks in LD. (G) Close-up view of the inflorescence of *dpa4-2 sod7-2 cuc2g-m4* mutant on CaAM. The plants were grown for 6 weeks in long-day-conditions. Arrows point to delayed CaAMs (H-I) Quantification of the transcript level of *CUC3* and *CUC2* by RT-qPCR in CaAM of wild-type plants and *dpa4-2 sod7-2* double mutant grown for 5 weeks in LD. Expressions were normalized using the QREF and REFA genes. A Student’s test was performed to compare the expression levels of the mutants in comparison to the wild-type (p <0.05 *; p <0.01 **). Scales bars: (A-D; F-G) = 5 cm

**Figure 4 Supplemental. Delayed development of *dpa4-2 sod7-2* is not restored by mutations in *BRC1*/*MAX* genes.** (A-H) Inflorescence of WT, single *max-brc1* mutants and *dpa4-2 sod7-2*-*max/brc1* triple mutants. Plants were grown for 5 weeks in LD. White arrowheads point to the developed CaAMs while arrows point to delayed CaAMs. The time point after bolting at which half of the CaAM are developed (t_50_) is indicated under the genotype. (I-K) Kinetics of CaAM development of single *max-brc1* mutants and *dpa4-2 sod7-2*-*max/brc1* triple mutant after bolting. Development of the CaAM is indicated as the percentage of developed branches (≥ 3mm) reported to the total number of cauline leaves (*n*≥8). All data were generated in the same experiments, therefore the same WT and *dpa4-2 sod7-2* data were used in panels I to K. Scales bars: (A-H) = 5 cm

**Figure 5 Supplemental. *DPA4* and *SOD7* have overlapping expression patterns with *CUC2* and *CUC3* in the boundary domain and are transiently expressed in the early AM** (A-H) Maximum projections of transverse optical sections of plant co-expressing pCUC3:erCFP and pSOD7:GFP reporters. (I-P) Maximum projections of transverse optical sections of plant co-expressing pCUC2:erCFP and pDPA4:GFP reporters CaAMs are at the (A,E,I,M) line, (B,F,J,N) eye, (C,G,K,O) dome and late dome stage (D,H,L,P) Scale bars: (A-P) = 50µm; sam: shoot apical meristem; clp: cauline leaf primordium; *: AM; the dotted line corresponds to the outline of the cauline leaf primordium.

**Figure 6 Supplemental. Putative NGAL binding sites in *CUC3* and pCUC3/pCUC3-6m reporter expression in CaAM.** (A) Diagram of *CUC3* promoter and CDS with all the putative *NGAL* binding sites identified (Swaminathan et al., 2008; Zhang et al., 2015). A focus on the sequence of BS1 in shown. A and A-m indicate the wild-type probe and the mutated probe used in the EMSA, respectively. (B) EMSA experiments showed that SOD7 directly binds to the promoter of CUC3. The biotin-labeled probe A and MBP-SOD7 formed a DNA-protein complex (lane 2), but the mutated probe A-m and MBP-SOD7 did not (lane 9). The biotin-labeled probe A and MBP did not form a DNA-protein complex (lane 1). The retarded DNA-protein complex was reduced by the competition using the unlabeled probe A (lane 3 to 5), but not reduced by the competition using the unlabeled mutated probe A-m (lane 6 to 8). (C-D) Maximum projections of transverse optical sections of pCUC3:mCherry-N7 or pCUC3-6m:GFP-N7 reporters in wild-type plants during CaAM formation at leaf primordium stage. Scale bars: (B-C) = 50µm; the dotted line corresponds to the outline of the cauline meristem and leaf primordia.

**Figure 7 Supplemental. *CLV3* and *WUS* expression patterns in CaAMs.** (A-D) Maximum projections of tangential optical sections of whole mount *in situ hybridization* of *CLV3* transcript in WT and *dpa4-2 sod7-2* in CaAMs at dome stage (E-F) and leaf primordium stage (G-H). (E,F) Maximum projections of tangential optical sections of the pCLV3:mCHERRY-NLS reporter during CaAM development at dome stage. (G-I) Maximum projections of optical sections of whole mount *in situ hybridization* of *WUS* transcript in CaAMs at dome stage. Scale bars: 50µm

**Figure 8 Supplemental. *WUS* and *CLV3* activation are delayed in *dpa4-2 sod7-2* floral meristems** (A,B) *WUS* and *CLV3* expression as a function of floral meristem size

