## Supplemental Figures and Tables for "*De novo* stem cell establishment in meristems requires repression of organ boundary cell fate"

Nicolas et al., Figure 1 Supplemental

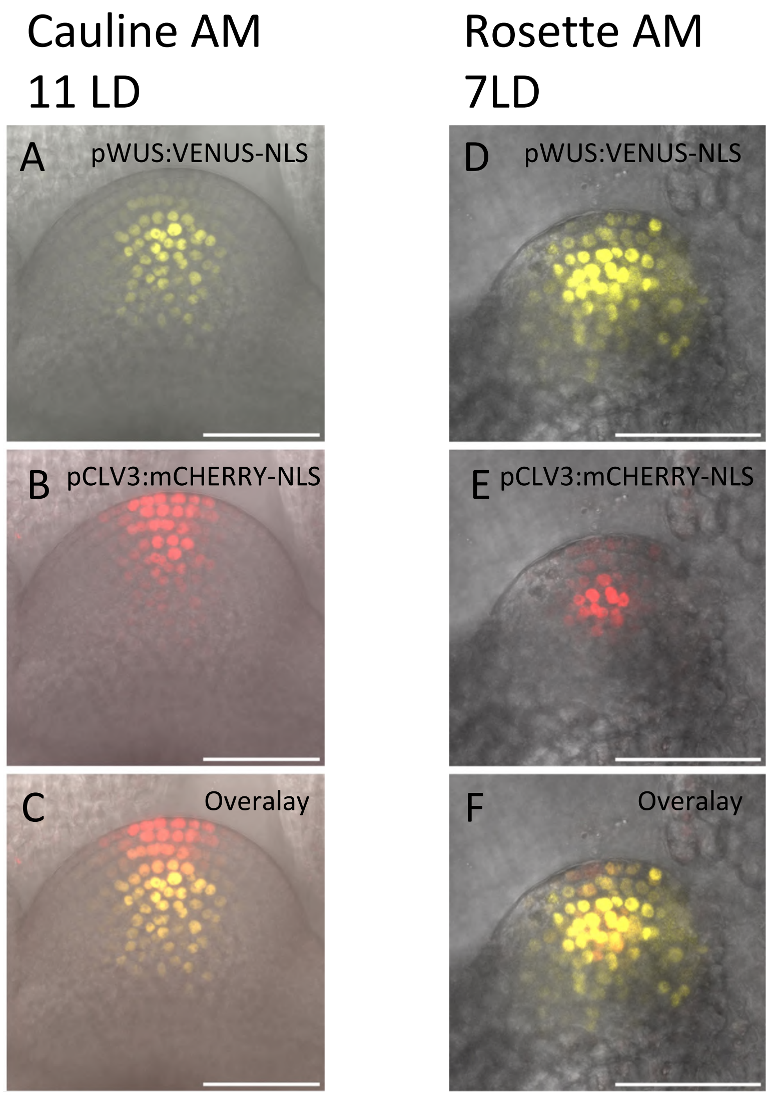


**Figure 1 Supplemental. *WUS* and *CLV3* expression in CaAM and RoAM**
(A-C) Maximum projections of radial optical sections of a pWUS:VENUS-NLS (A) and pCLV3:mCHERRY-NLS (B) reporter lines and the overlay (C) during CaAM formation.
(D-F) Maximum projections of radial optical sections of a pWUS:VENUS-NLS (D) and pCLV3:mCHERRY-NLS (E) reporter lines and the overlay (F) during RoAM formation.
The number of days in LD conditions is indicated.
Scale bars : (A-F) = 50 µm

### Nicolas et al., Figure 2 Supplemental

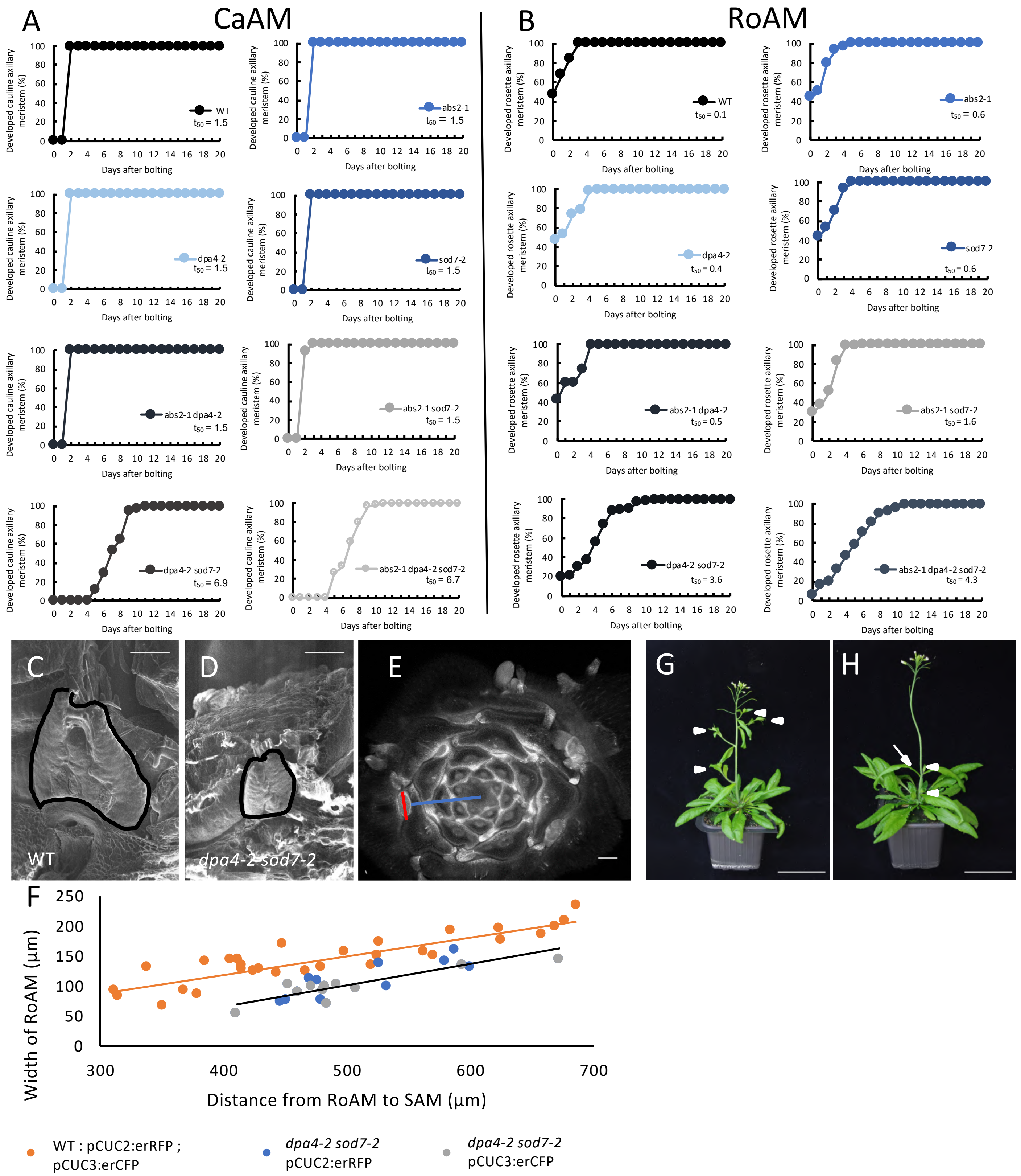


**Figure 2 Supplemental. *DPA4* and *SOD7* are required for rapid development of cauline AMs.**

(A,B) Kinetics of CaAM or RoAM development of all *ngal* simple and multiple mutants after bolting. Development of the meristems is indicated as the percentage of developed branches (≥ 3mm) reported to the total number of cauline or rosette leaves (*n*≥11).

(C,D) SEM observations of WT and *dpa4-2 sod7-2* RoAM from leaf 7 on plants grown 4 weeks in SD.

(E) Quantification method for (F) on a maximum projection of transverse optical sections of pCUC3:erCFP reporter in WT SAM on plants grown 4 weeks in SD. The red line represents the width of the RoAM and the blue line the distance between the SAM and the RoAM.

(F) RoAM width as a function of the distance between the same RoAM and the SAM .

(G,H) Inflorescence of WT and *dpa4-3 sod7-2* double mutants. Plants were grown for 5 weeks in LD. White arrowheads point to the developed CaAMs while the arrows point to delayed CaAMs.

Scale bars : (C,D) = 200 µm ; (E) = 100 µm, (G,H) = 5cm

### Nicolas et al., Figure 3 Supplemental

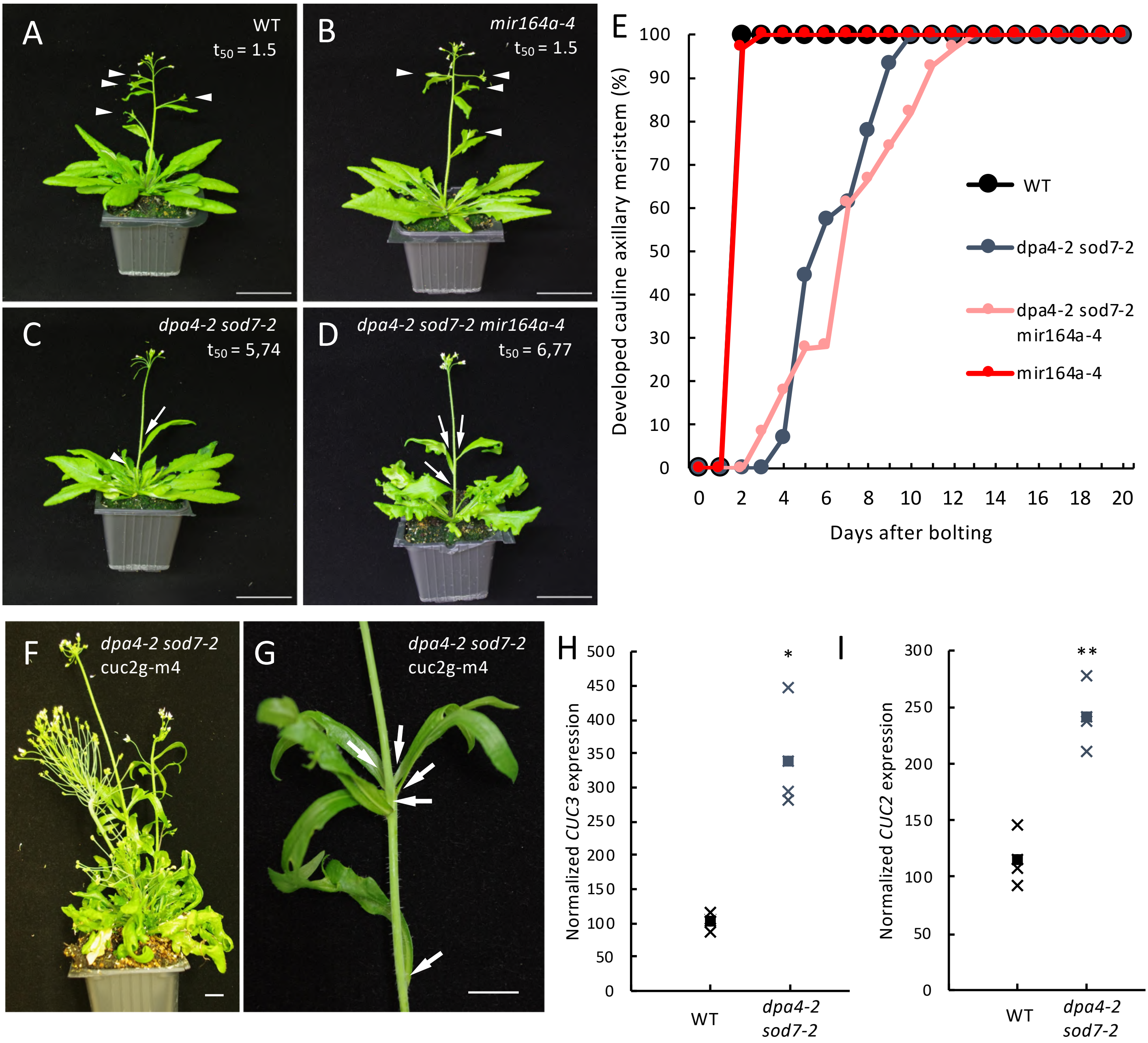


**Figure 3 Supplemental. Genetic interaction between *MIR164* and *DPA4/SOD7* during CaAM development and *CUC2/CUC3* mRNA quantification in *dpa4-2 sod7-2*.**

(A-D) Inflorescence of WT, and *mir164a-4*, *dpa4-2 sod7-2* and *dpa4-2 sod7-2 mir164a-4* mutants. Plants were grown for 5 weeks in LD. White arrowheads point to the developed CaAMs while the arrows point to delayed CaAMs. The time point after bolting at which half of the CaAM are developed ($t_{50}$) is indicated under the genotype.

(E) Kinetics of CaAM development after bolting. Development of the CaAM is indicated as the percentage of developed branches (≥ 3mm) reported to the total number of cauline leaves ($n \geq 8$).

(F) Inflorescence of *dpa4-2 sod7-2 cuc2g-m4* mutant. Plants were grown for 6 weeks in LD.

(G) Close-up view of the inflorescence of *dpa4-2 sod7-2 cuc2g-m4* mutant on CaAM. The plants were grown for 6 weeks in long-day-conditions. Arrows point to delayed CaAMs

(H-I) Quantification of the transcript level of *CUC3* and *CUC2* by RT-qPCR in CaAM of wild-type plants and *dpa4-2 sod7-2* double mutant grown for 5 weeks in LD. Expressions were normalized using the QREF and REFA genes. A Student's test was performed to compare the expression levels of the mutants in comparison to the wild-type ($p < 0.05$ *; $p < 0.01$ **).

Scales bars : (A-D ; F-G) = 5 cm

Nicolas et al., Figure 4 Supplemental

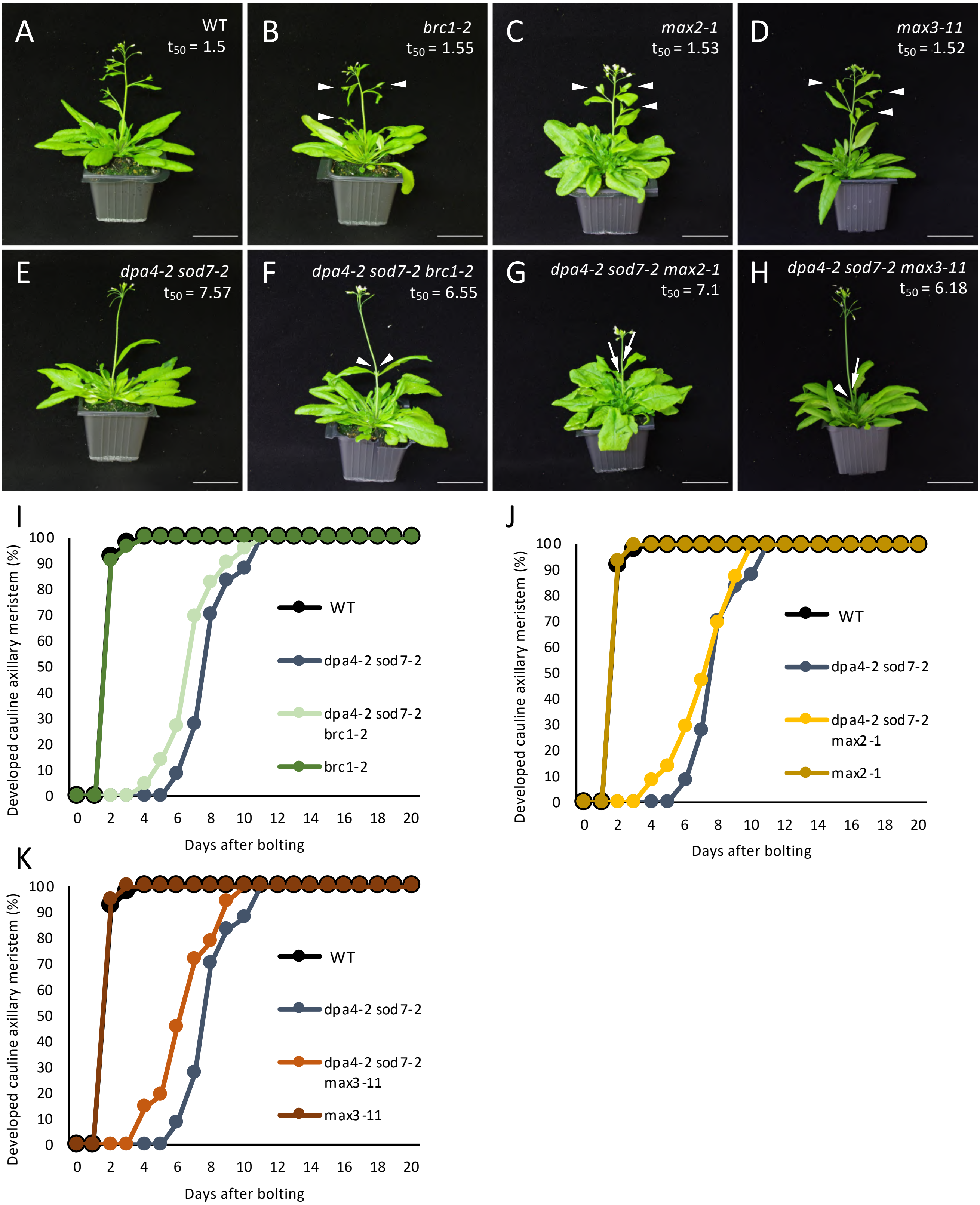


**Figure 4 Supplemental. Delayed development of *dpa4-2 sod7-2* is not restored by mutations in *BRC1/MAX* genes.**

(A-H) Inflorescence of WT, single *max-brc1* mutants and *dpa4-2 sod7-2-max/brc1* triple mutants. Plants were grown for 5 weeks in LD. White arrowheads point to the developed CaAMs while arrows point to delayed CaAMs. The time point after bolting at which half of the CaAM are developed ($t_{50}$) is indicated under the genotype.

(I-K) Kinetics of CaAM development of single *max-brc1* mutants and *dpa4-2 sod7-2-max/brc1* triple mutant after bolting. Development of the CaAM is indicated as the percentage of developed branches (≥ 3mm) reported to the total number of cauline leaves (*n*≥8). All data were generated in the same experiments, therefore the same WT and *dpa4-2 sod7-2* data were used in panels I to K.

Scales bars : (A-H) = 5 cm

### Nicolas et al., Figure 5 Supplemental

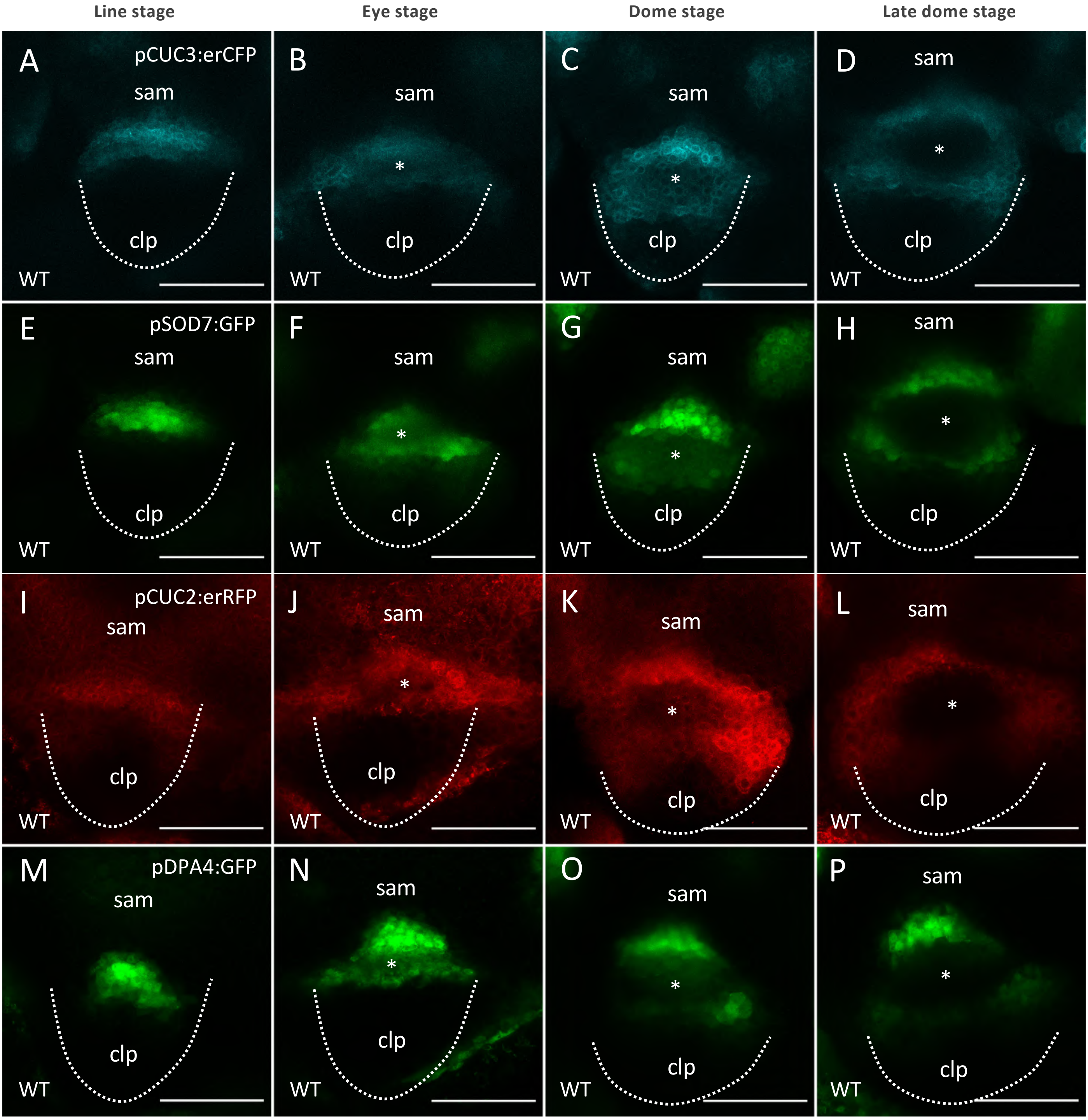


**Figure 5 Supplemental. *DPA4* and *SOD7* have overlapping expression patterns with *CUC2* and *CUC3* in the boundary domain and are transiently expressed in the early AM**

(A-H) Maximum projections of transverse optical sections of plant co-expressing pCUC3:erCFP and pSOD7:GFP reporters.

(I-P) Maximum projections of transverse optical sections of plant co-expressing pCUC2:erCFP and pDPA4:GFP reporters

CaAMs are at the (A,E,I,M) line, (B,F,J,N) eye, (C,G,K,O) dome and late dome stage (D,H,L,P)

Scale bars : (A-P) = 50µm; sam: shoot apical meristem; clp: cauline leaf primordium; *: AM ; the dotted line corresponds to the outline of the cauline leaf primordium.

### Nicolas et al., Figure 6 Supplemental

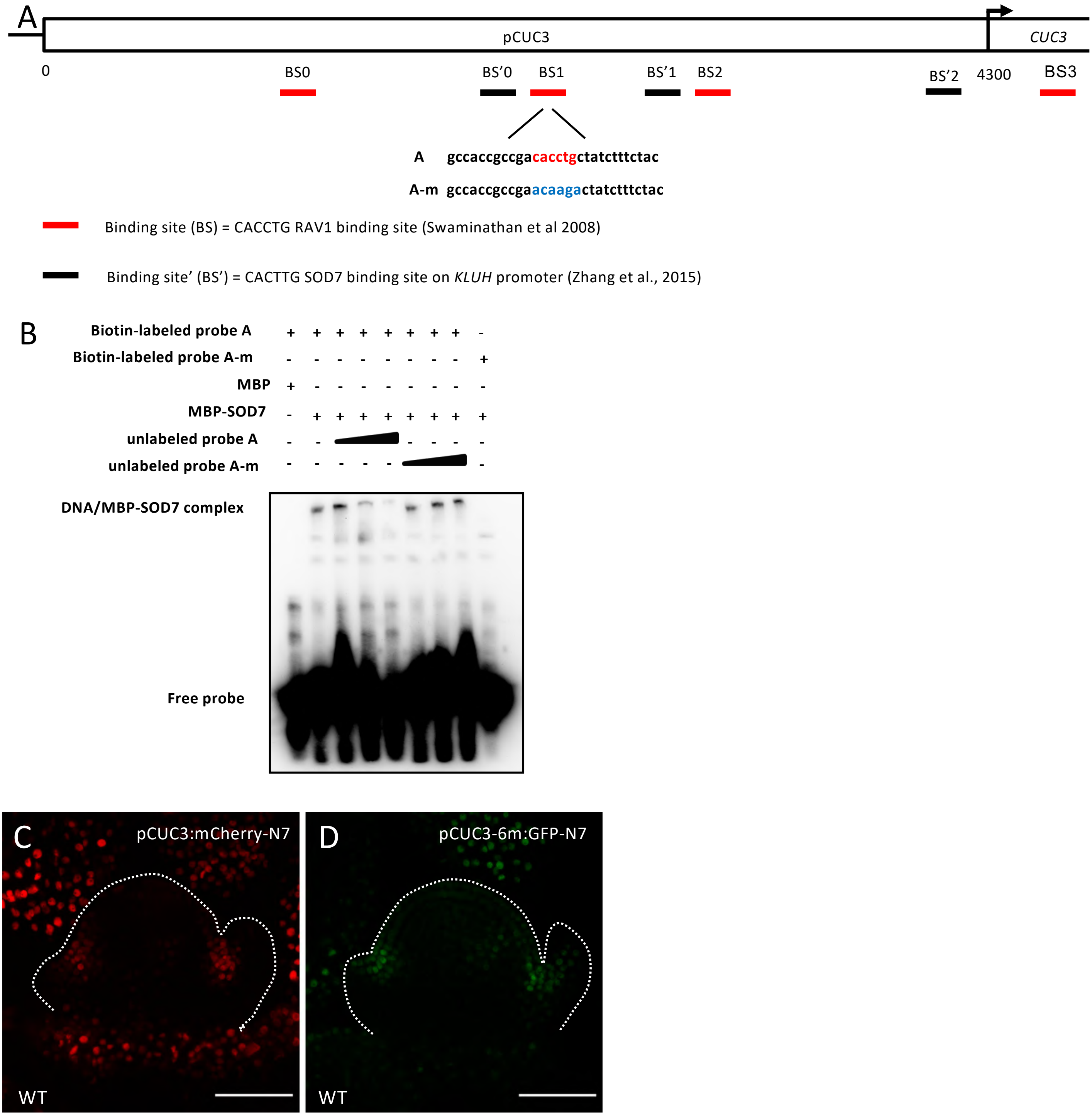


**Figure 6 Supplemental. Putative NGAL binding sites in *CUC3* and pCUC3/pCUC3-6m reporter expression in CaAM.**

(A) Diagram of *CUC3* promoter and CDS with all the putative *NGAL* binding sites identified (Swaminathan et al., 2008 ; Zhang et al., 2015).

A focus on the sequence of BS1 in shown. A and A-m indicate the wild-type probe and the mutated probe used in the EMSA, respectively.

(B) EMSA experiments showed that SOD7 directly binds to the promoter of CUC3. The biotin-labeled probe A and MBP-SOD7 formed a DNA-protein complex (lane 2), but the mutated probe A-m and MBP-SOD7 did not (lane 9). The biotin-labeled probe A and MBP did not form a DNA-protein complex (lane 1). The retarded DNA-protein complex was reduced by the competition using the unlabeled probe A (lane 3 to 5), but not reduced by the competition using the unlabeled mutated probe A-m (lane 6 to 8).

(C-D) Maximum projections of transverse optical sections of pCUC3:mCherry-N7 or pCUC3-6m:GFP-N7 reporters in wild-type plants during CaAM formation at leaf primordium stage.

Scale bars : (B-C) = 50µm ; the dotted line corresponds to the outline of the cauline meristem and leaf primordia.

Nicolas et al., Figure 7 Supplemental

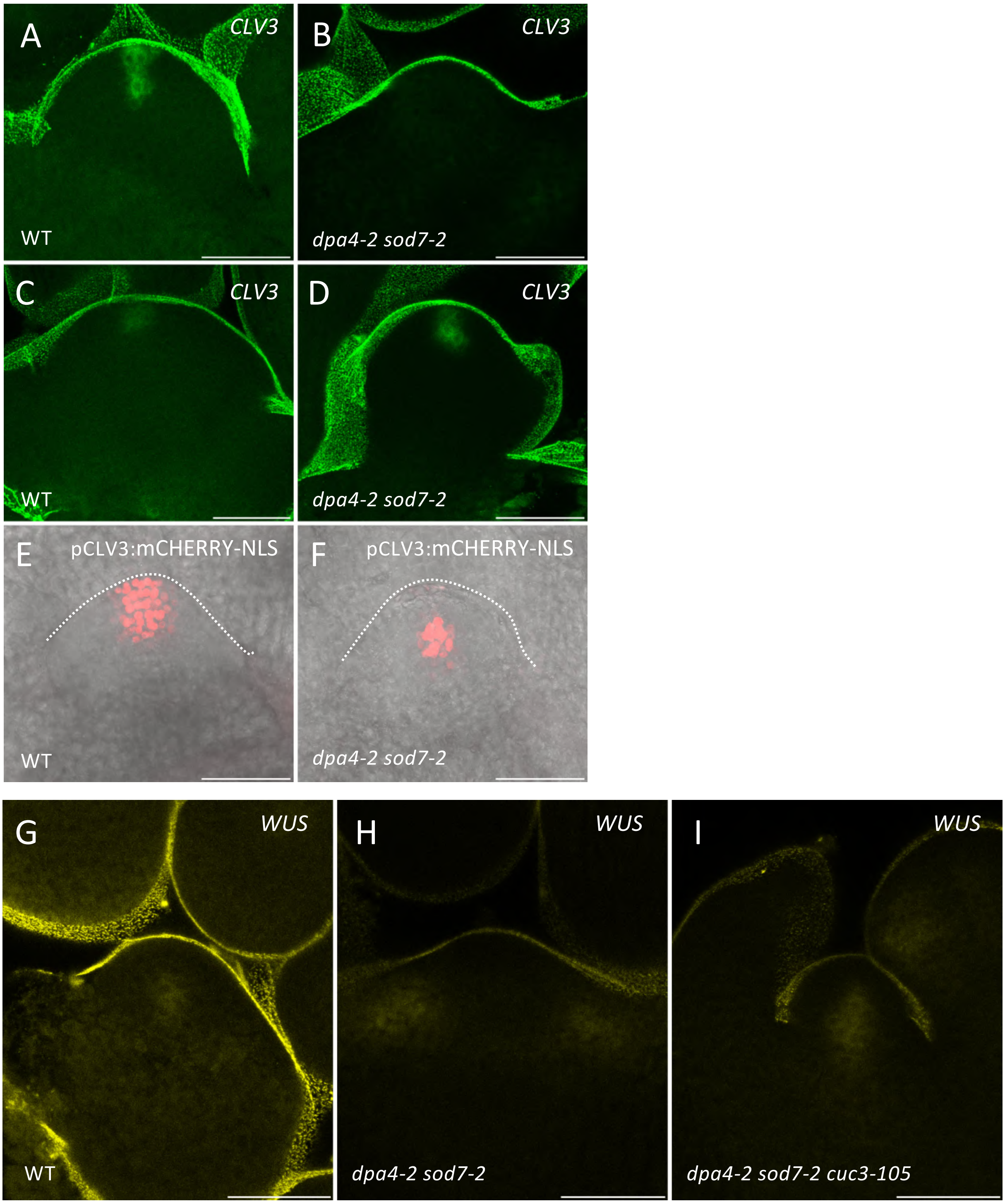


**Figure 7 Supplemental. *CLV3* and *WUS* expression patterns in CaAMs.**
(A-D) Maximum projections of tangential optical sections of whole mount *in situ hybridization* of *CLV3* transcript in WT and *dpa4-2 sod7-2* in CaAMs at dome stage (E-F) and leaf primordium stage (G-H).
(E,F) Maximum projections of tangential optical sections of the pCLV3:mCHERRY-NLS reporter during CaAM development at dome stage.
(G-I) Maximum projections of optical sections of whole mount *in situ hybridization* of *WUS* transcript in CaAMs at dome stage.
Scale bars : 50µm

### Nicolas et al., Figure 8 Supplemental

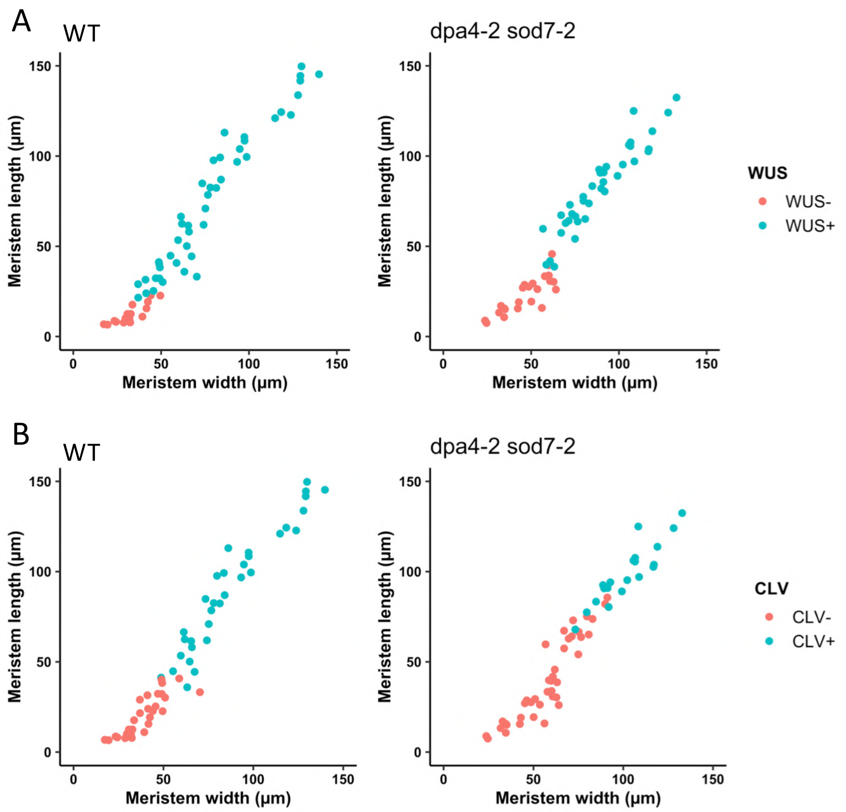


**Figure 8 Supplemental. *WUS* and *CLV3* activation are delayed in *dpa4-2 sod7-2* floral meristems**
(A,B) *WUS* and *CLV3* expression as a function of floral meristem size

**Supplemental Table 1. Primers used in this work**

| Primer Name | Sequence (5'-3') | Aim | Reference |
|---|---|---|---|
| Pdpa4-2FW | GGGGACAAGTTTGTACAAAAAAGCAGGCTAAATCACGGGACGGATAAAGA | Cloning pDPA4 | |
| Pdpa4-2RV | GGGGACCACTTTGTACAAGAAAGCTGGGTGTTAGGTGAAAGAGAGAGGTGGG | Cloning pDPA4 | |
| SOD7Profwattb1 | GGGGACAAGTTTGTACAAAAAAGCAGGCTAAACACGTCAAATATAACGAAT | Cloning pSOD7 | |
| SOD7Prorvattb2 | GGGGACCACTTTGTACAAGAAAGCTGGGTCTTTTTTTTTGGTTTCTTGGAGTGAGAGAGAGAG | Cloning pSOD7 | |
| GB_S1pCUC3S2_F | GCGCCGTCTCGCTCGGGAGCGGCACTGGATACACGAACA | Cloning pCUC3 | |
| GB_S1pCUC3S2_R | GCGCCGTCTCGCTCACATTGAAGAACAAAGATAGCAATGATAC | Cloning pCUC3 | |
| promCUC2 Fwd | GATATCATCAAGGCATATTTAAA | Cloning of pCUC2 for Y1H | |
| promCUC2 Rv | AGATCTAAAGCTTTTGTTTGAGA | Cloning of pCUC2 for Y1H | |
| prCuc3 - Fw | CGGTCGACCGGCACTGGATACACG | Cloning of pCUC3 for Y1H | |
| prCuc3 – R | CGCTGCAGGAAGAACAAAGATAGC | Cloning of pCUC3 for Y1H | |
| CUC3_S2F | GCGCCGTCTCGCTCGAATGATGCTTGCGGTGGAAGA | Cloning patch1 of *CUC3* CDS and patch1 of *CUC3-1m* CDS | |
| CUC3_dom1R | GCGCCGTCTCGAAAGACCACCATGGAAGATTTTG | Cloning patch1 of *CUC3* CDS | |
| CUC3_dom1F | GCGCCGTCTCGCTTTCCGGCATTCACATTTCC | Cloning patch2 of *CUC3* CDS | |
| CUC3_dom2R | GCGCCGTCTCGATACGACAGGGTTGATTAAGG | Cloning patch2 of *CUC3* CDS | |
| CUC3_dom2F | GCGCCGTCTCGGTATCTCAGCTCAACGGTATC | Cloning patch3 of *CUC3* CDS | |
| CUC3_S7R | GCGCCGTCTCGCTCAAAGCCTACAGCTGGAATCCTAAAGG | Cloning patch3 of *CUC3* CDS and patch2 of *CUC3-1m* CDS | |
| CUC3_CDS_PF3_f | GCGCCGTCTCTTGCCGCCGGGCTTCCGGTTTC | Cloning patch1 of *CUC3-1m* CDS | |
| CUC3_CDS_PF3_r | GCGCCGTCTCGCAATCCTCTCTCGTTCCTTTC | Cloning patch2 of *CUC3-1m* CDS | |
| qPCR_CUC3_3_f | CTCCGGCATTCACATTTCCGA | qRT-PCR | |
| qPCR_CUC3_3_r | AGCTTTCCAGTATCCAGCAGT | qRT-PCR | |
| qREF-F | AACTCTATGCAGCATTTGATCCACT | qRT-PCR | Morineau, et al., 2016 |
| qREF-R | TGATTGCATATCTTTATCGCCATC | qRT-PCR | Morineau, et al., 2016 |
| qCUC2-F | CTTGGCAACTTCCCGGGAGA | qRT-PCR | Tian, et al., 2014 |
| qCUC2-R | CCAGCCTCAGTTGCTCTGTTAGTT | qRT-PCR | Tian, et al., 2014 |
| REFA_F | GAGCTGAAGTGGCTTCCATGAC | qRT-PCR | Czechowski et al., 2005 |
| REFA_R | GGTCCGACATACCCATGATCC | qRT-PCR | Czechowski et al., 2005 |
| insituCUC3 L Fwd | ATGATGCTTGCGGTGGAAGA | *in situ* | |
| insituCUC3 L Rv-T | TGTAATACGACTCACTATAGGGCCTACAGCTGGAATCCTAAA | *in situ* | |
| insituCUC2 L Fwd | ATGGACATTCCGTATTACCA | *in situ* | |
| insitu CUC2 Rv-T | TGTAATACGACTCACTATAGGGCTCAGTAGTTCCAAATACA | *in situ* | |
| ISH WUS b AS T7 | TGTAATACGACTCACTATAGGGCCCATCCTCCACCTACGTTGT | in situ | |
| ISH WUSb Sens R | CCATCCTCCACCTACGTTGT | in situ | |
| AS-CLV3-4L | AAAAATGGATTCGAAGAGTTTTCTGC | in situ | |
| AS-CLV3-4R-T7 | TGTAATACGACTCACTATAGGGCAAGAGATTAGGTCAAGGGAGCTGA | in situ | |


Czechowski, T., Stitt, M., Altmann, T., Udvardi, M.K., and Scheible, W.-R. (2005). Genome-Wide Identification and Testing of Superior Reference Genes for Transcript Normalization in Arabidopsis. Plant Physiol. 139, 5–17.

Morineau, C., Gissot, L., Bellec, Y., Hematy, K., Tellier, F., Renne, C., Haslam, R., Beaudoin, F., Napier, J., and Faure, J.D. (2016). Dual fatty acid elongase complex interactions in arabidopsis. PLoS One *1*

Tian, C., Zhang, X., He, J., Yu, H., Wang, Y., Shi, B., Han, Y., Wang, G., Feng, X., Zhang, C., et al. (2014). An organ boundary-enriched gene regulatory network uncovers regulatory hierarchies underlying axillary meristem initiation. Mol. Syst. Biol. 10, 755–755.

**Supplemental Table 2. Prey transcriptio**

| AGI number | TF Family |
|---|---|
| AT2G36080 | ABI3/VP1 |
| AT3G11580 | ABI3/VP1 |
| AT3G18990 | ABI3/VP1 |
| AT4G32010 | ABI3/VP1 |
| AT1G14510 | Alfin-like |
| AT2G02470 | Alfin-like |
| AT3G11200 | Alfin-like |
| AT3G42790 | Alfin-like |
| AT5G05610 | Alfin-like |
| AT5G26210 | Alfin-like |
| AT1G12610 | AP2/EREBP |
| AT1G12630 | AP2/EREBP |
| AT1G12980 | AP2/EREBP |
| AT1G13260 | AP2/EREBP |
| AT1G15360 | AP2/EREBP |
| AT1G19210 | AP2/EREBP |
| AT1G21910 | AP2/EREBP |
| AT1G22190 | AP2/EREBP |
| AT1G22810 | AP2/EREBP |
| AT1G22985 | AP2/EREBP |
| AT1G24590 | AP2/EREBP |
| AT1G25470 | AP2/EREBP |
| AT1G25560 | AP2/EREBP |
| AT1G28360 | AP2/EREBP |
| AT1G28370 | AP2/EREBP |
| AT1G36060 | AP2/EREBP |
| AT1G43160 | AP2/EREBP |
| AT1G44830 | AP2/EREBP |
| AT1G50640 | AP2/EREBP |
| AT1G51190 | AP2/EREBP |
| AT1G53170 | AP2/EREBP |
| AT1G53910 | AP2/EREBP |
| AT1G64380 | AP2/EREBP |
| AT1G68550 | AP2/EREBP |
| AT1G68840 | AP2/EREBP |
| AT1G71130 | AP2/EREBP |
| AT1G71450 | AP2/EREBP |
| AT1G72360 | AP2/EREBP |
| AT1G74930 | AP2/EREBP |
| AT1G77640 | AP2/EREBP |
| AT1G78080 | AP2/EREBP |
| AT1G79700 | AP2/EREBP |
| AT2G20880 | AP2/EREBP |
| AT2G23340 | AP2/EREBP |
| AT2G28550 | AP2/EREBP |
| AT2G31230 | AP2/EREBP |
| AT2G38340 | AP2/EREBP |
| AT2G41710 | AP2/EREBP |
| AT2G44840 | AP2/EREBP |
| AT2G44940 | AP2/EREBP |
| AT2G46310 | AP2/EREBP |
| AT2G47520 | AP2/EREBP |
| AT3G11020 | AP2/EREBP |
| AT3G14230 | AP2/EREBP |
| AT3G15210 | AP2/EREBP |
| AT3G16280 | AP2/EREBP |
| AT3G16770 | AP2/EREBP |
| AT3G20310 | AP2/EREBP |
| AT3G20840 | AP2/EREBP |
| AT3G23240 | AP2/EREBP |
| AT3G25730 | AP2/EREBP |
| AT3G25890 | AP2/EREBP |
| AT3G50260 | AP2/EREBP |
| AT3G54990 | AP2/EREBP |

| | |
|---|---|
| AT3G60490 | AP2/EREBP |
| AT3G61630 | AP2/EREBP |
| AT4G11140 | AP2/EREBP |
| AT4G16750 | AP2/EREBP |
| AT4G17490 | AP2/EREBP |
| AT4G17500 | AP2/EREBP |
| AT4G18450 | AP2/EREBP |
| AT4G23750 | AP2/EREBP |
| AT4G25470 | AP2/EREBP |
| AT4G25490 | AP2/EREBP |
| AT4G27950 | AP2/EREBP |
| AT4G28140 | AP2/EREBP |
| AT4G32800 | AP2/EREBP |
| AT4G34410 | AP2/EREBP |
| AT4G36900 | AP2/EREBP |
| AT4G36920 | AP2/EREBP |
| AT4G37750 | AP2/EREBP |
| AT4G39780 | AP2/EREBP |
| AT5G05410 | AP2/EREBP |
| AT5G07580 | AP2/EREBP |
| AT5G10510 | AP2/EREBP |
| AT5G13330 | AP2/EREBP |
| AT5G13910 | AP2/EREBP |
| AT5G17430 | AP2/EREBP |
| AT5G18560 | AP2/EREBP |
| AT5G25390 | AP2/EREBP |
| AT5G25810 | AP2/EREBP |
| AT5G44210 | AP2/EREBP |
| AT5G47220 | AP2/EREBP |
| AT5G47230 | AP2/EREBP |
| AT5G51190 | AP2/EREBP |
| AT5G51990 | AP2/EREBP |
| AT5G52020 | AP2/EREBP |
| AT5G53290 | AP2/EREBP |
| AT5G57390 | AP2/EREBP |
| AT5G60120 | AP2/EREBP |
| AT5G61590 | AP2/EREBP |
| AT5G61600 | AP2/EREBP |
| AT5G61890 | AP2/EREBP |
| AT5G65510 | AP2/EREBP |
| AT5G67190 | AP2/EREBP |
| AT1G19220 | ARF |
| AT1G19850 | ARF |
| AT1G30330 | ARF |
| AT1G34170 | ARF |
| AT1G34310 | ARF |
| AT1G34390 | ARF |
| AT1G35240 | ARF |
| AT1G35540 | ARF |
| AT1G43950 | ARF |
| AT1G59750 | ARF |
| AT1G77850 | ARF |
| AT2G28350 | ARF |
| AT2G33860 | ARF |
| AT2G46530 | ARF |
| AT3G61830 | ARF |
| AT4G23980 | ARF |
| AT4G30080 | ARF |
| AT5G20730 | ARF |
| AT5G20730 | ARF |
| AT5G37020 | ARF |
| AT5G60450 | ARF |
| AT5G62000 | ARF |
| AT1G04880 | ARID |
| AT1G20910 | ARID |
| AT1G76510 | ARID |
| AT3G43240 | ARID |
| AT1G16530 | AS2 |

| | |
|---|---|
| AT1G31320 | AS2 |
| AT3G49940 | AS2 |
| AT4G00220 | AS2 |
| AT4G37540 | AS2 |
| AT5G63090 | AS2 |
| AT5G67420 | AS2 |
| AT1G04100 | AUX/IAA |
| AT1G04240 | AUX/IAA |
| AT1G04250 | AUX/IAA |
| AT1G04550 | AUX/IAA |
| AT1G15580 | AUX/IAA |
| AT1G51950 | AUX/IAA |
| AT1G52830 | AUX/IAA |
| AT2G22670 | AUX/IAA |
| AT2G33310 | AUX/IAA |
| AT2G46990 | AUX/IAA |
| AT3G04730 | AUX/IAA |
| AT3G15540 | AUX/IAA |
| AT3G16500 | AUX/IAA |
| AT3G17600 | AUX/IAA |
| AT3G23030 | AUX/IAA |
| AT3G23050 | AUX/IAA |
| AT3G62100 | AUX/IAA |
| AT4G14550 | AUX/IAA |
| AT4G14560 | AUX/IAA |
| AT4G28640 | AUX/IAA |
| AT4G29080 | AUX/IAA |
| AT5G25890 | AUX/IAA |
| AT5G43700 | AUX/IAA |
| AT5G65670 | AUX/IAA |
| AT1G19350 | BES1 |
| AT1G75080 | BES1 |
| AT1G01260 | bHLH |
| AT1G02340 | bHLH |
| AT1G03040 | bHLH |
| AT1G05710 | bHLH |
| AT1G05805 | bHLH |
| AT1G09250 | bHLH |
| AT1G09530 | bHLH |
| AT1G10120 | bHLH |
| AT1G12860 | bHLH |
| AT1G22490 | bHLH |
| AT1G31050 | bHLH |
| AT1G32640 | bHLH |
| AT1G51070 | bHLH |
| AT1G51140 | bHLH |
| AT1G59640 | bHLH |
| AT1G61660 | bHLH |
| AT1G68810 | bHLH |
| AT1G68920 | bHLH |
| AT1G69010 | bHLH |
| AT1G74500 | bHLH |
| AT2G16910 | bHLH |
| AT2G20180 | bHLH |
| AT2G22760 | bHLH |
| AT2G22770 | bHLH |
| AT2G24260 | bHLH |
| AT2G31730 | bHLH |
| AT2G43010 | bHLH |
| AT2G43060 | bHLH |
| AT2G46510 | bHLH |
| AT2G47270 | bHLH |
| AT3G05800 | bHLH |
| AT3G06120 | bHLH |
| AT3G06590 | bHLH |
| AT3G07340 | bHLH |
| AT3G17100 | bHLH |
| AT3G19860 | bHLH |

| | |
|---|---|
| AT3G23210 | bHLH |
| AT3G23690 | bHLH |
| AT3G24140 | bHLH |
| AT3G25710 | bHLH |
| AT3G26744 | bHLH |
| AT3G47640 | bHLH |
| AT3G56980 | bHLH |
| AT3G57800 | bHLH |
| AT3G59060 | bHLH |
| AT3G62090 | bHLH |
| AT4G02590 | bHLH |
| AT4G05170 | bHLH |
| AT4G14410 | bHLH |
| AT4G16430 | bHLH |
| AT4G17880 | bHLH |
| AT4G21340 | bHLH |
| AT4G30410 | bHLH |
| AT4G36540 | bHLH |
| AT4G36930 | bHLH |
| AT4G37850 | bHLH |
| AT5G08130 | bHLH |
| AT5G41315 | bHLH |
| AT5G46760 | bHLH |
| AT5G48560 | bHLH |
| AT5G53210 | bHLH |
| AT5G54680 | bHLH |
| AT5G58010 | bHLH |
| AT5G62610 | bHLH |
| AT5G65640 | bHLH |
| AT5G67110 | bHLH |
| AT1G03970 | bZIP |
| AT1G06070 | bZIP |
| AT1G08320 | bZIP |
| AT1G22070 | bZIP |
| AT1G32150 | bZIP |
| AT1G42990 | bZIP |
| AT1G43700 | bZIP |
| AT1G45249 | bZIP |
| AT1G49720 | bZIP |
| AT1G68640 | bZIP |
| AT1G77920 | bZIP |
| AT2G21230 | bZIP |
| AT2G22850 | bZIP |
| AT2G31370 | bZIP |
| AT2G35530 | bZIP |
| AT2G36270 | bZIP |
| AT2G40620 | bZIP |
| AT2G40950 | bZIP |
| AT2G41070 | bZIP |
| AT2G46270 | bZIP |
| AT3G12250 | bZIP |
| AT3G17609 | bZIP |
| AT3G19290 | bZIP |
| AT3G49760 | bZIP |
| AT3G54620 | bZIP |
| AT3G56850 | bZIP |
| AT3G58120 | bZIP |
| AT4G01120 | bZIP |
| AT4G02640 | bZIP |
| AT4G34000 | bZIP |
| AT4G35040 | bZIP |
| AT4G35900 | bZIP |
| AT4G36730 | bZIP |
| AT4G37730 | bZIP |
| AT4G38900 | bZIP |
| AT5G06839 | bZIP |
| AT5G06950 | bZIP |
| AT5G06960 | bZIP |

| | |
|---|---|
| AT5G10030 | bZIP |
| AT5G11260 | bZIP |
| AT5G24800 | bZIP |
| AT5G28770 | bZIP |
| AT5G44080 | bZIP |
| AT5G49450 | bZIP |
| AT5G65210 | bZIP |
| AT1G73870 | C2C2-CO-like |
| AT1G75540 | C2C2-CO-like |
| AT1G78600 | C2C2-CO-like |
| AT2G21320 | C2C2-CO-like |
| AT2G24790 | C2C2-CO-like |
| AT2G47890 | C2C2-CO-like |
| AT3G07650 | C2C2-CO-like |
| AT3G21890 | C2C2-CO-like |
| AT4G38960 | C2C2-CO-like |
| AT4G39070 | C2C2-CO-like |
| AT5G15840 | C2C2-CO-like |
| AT5G24930 | C2C2-CO-like |
| AT5G48250 | C2C2-CO-like |
| AT5G54470 | C2C2-CO-like |
| AT5G57660 | C2C2-CO-like |
| AT5G61380 | C2C2-CO-like |
| AT1G07640 | C2C2-Dof |
| AT1G29160 | C2C2-Dof |
| AT1G51700 | C2C2-Dof |
| AT1G64620 | C2C2-Dof |
| AT2G28510 | C2C2-Dof |
| AT2G28810 | C2C2-Dof |
| AT2G34140 | C2C2-Dof |
| AT2G37590 | C2C2-Dof |
| AT2G46590 | C2C2-Dof |
| AT3G21270 | C2C2-Dof |
| AT3G45610 | C2C2-Dof |
| AT3G47500 | C2C2-Dof |
| AT3G50410 | C2C2-Dof |
| AT3G55370 | C2C2-Dof |
| AT3G61850 | C2C2-Dof |
| AT4G00940 | C2C2-Dof |
| AT4G24060 | C2C2-Dof |
| AT5G02460 | C2C2-Dof |
| AT5G39660 | C2C2-Dof |
| AT5G60200 | C2C2-Dof |
| AT5G60850 | C2C2-Dof |
| AT5G62430 | C2C2-Dof |
| AT5G62940 | C2C2-Dof |
| AT1G51600 | C2C2-GATA |
| AT2G18380 | C2C2-GATA |
| AT3G06740 | C2C2-GATA |
| AT3G16870 | C2C2-GATA |
| AT3G21175 | C2C2-GATA |
| AT3G24050 | C2C2-GATA |
| AT3G54810 | C2C2-GATA |
| AT3G60530 | C2C2-GATA |
| AT4G17570 | C2C2-GATA |
| AT4G24470 | C2C2-GATA |
| AT4G32890 | C2C2-GATA |
| AT4G34680 | C2C2-GATA |
| AT4G36620 | C2C2-GATA |
| AT5G25830 | C2C2-GATA |
| AT5G26930 | C2C2-GATA |
| AT5G56860 | C2C2-GATA |
| AT5G66320 | C2C2-GATA |
| AT1G08465 | C2C2-YABBY |
| AT1G69180 | C2C2-YABBY |
| AT2G26580 | C2C2-YABBY |
| AT2G45190 | C2C2-YABBY |
| AT4G00180 | C2C2-YABBY |

| | |
|---|---|
| AT1G03840 | C2H2 |
| AT1G04850 | C2H2 |
| AT1G10480 | C2H2 |
| AT1G14580 | C2H2 |
| AT1G24625 | C2H2 |
| AT1G26610 | C2H2 |
| AT1G27730 | C2H2 |
| AT1G30970 | C2H2 |
| AT1G34370 | C2H2 |
| AT1G43860 | C2H2 |
| AT1G50670 | C2H2 |
| AT1G55110 | C2H2 |
| AT1G55460 | C2H2 |
| AT1G66140 | C2H2 |
| AT1G67030 | C2H2 |
| AT1G68130 | C2H2 |
| AT1G68360 | C2H2 |
| AT1G68480 | C2H2 |
| AT1G72050 | C2H2 |
| AT1G75710 | C2H2 |
| AT2G01650 | C2H2 |
| AT2G01940 | C2H2 |
| AT2G02070 | C2H2 |
| AT2G19380 | C2H2 |
| AT2G24500 | C2H2 |
| AT2G26940 | C2H2 |
| AT2G27100 | C2H2 |
| AT2G28200 | C2H2 |
| AT2G29660 | C2H2 |
| AT2G36930 | C2H2 |
| AT2G37430 | C2H2 |
| AT2G41940 | C2H2 |
| AT2G45120 | C2H2 |
| AT2G48100 | C2H2 |
| AT3G02790 | C2H2 |
| AT3G02860 | C2H2 |
| AT3G05760 | C2H2 |
| AT3G10470 | C2H2 |
| AT3G12270 | C2H2 |
| AT3G13810 | C2H2 |
| AT3G19580 | C2H2 |
| AT3G23130 | C2H2 |
| AT3G44750 | C2H2 |
| AT3G49930 | C2H2 |
| AT3G53600 | C2H2 |
| AT3G57480 | C2H2 |
| AT3G58070 | C2H2 |
| AT3G60580 | C2H2 |
| AT3G62240 | C2H2 |
| AT4G16610 | C2H2 |
| AT4G16845 | C2H2 |
| AT4G27240 | C2H2 |
| AT4G31420 | C2H2 |
| AT5G01160 | C2H2 |
| AT5G03510 | C2H2 |
| AT5G03740 | C2H2 |
| AT5G04340 | C2H2 |
| AT5G04390 | C2H2 |
| AT5G14010 | C2H2 |
| AT5G16470 | C2H2 |
| AT5G25160 | C2H2 |
| AT5G26610 | C2H2 |
| AT5G40710 | C2H2 |
| AT5G43170 | C2H2 |
| AT5G44160 | C2H2 |
| AT5G51230 | C2H2 |
| AT5G52010 | C2H2 |
| AT5G59820 | C2H2 |

| | |
|---|---|
| AT5G63280 | C2H2 |
| AT5G66730 | C2H2 |
| AT1G68200 | C3H |
| AT5G08190 | CCAAT-Dr1 |
| AT5G23090 | CCAAT-Dr1 |
| AT1G54160 | CCAAT-HAP2 |
| AT1G72830 | CCAAT-HAP2 |
| AT5G12840 | CCAAT-HAP2 |
| AT2G38880 | CCAAT-HAP3 |
| AT3G53340 | CCAAT-HAP3 |
| AT5G47640 | CCAAT-HAP3 |
| AT1G07980 | CCAAT-HAP5 |
| AT1G08970 | CCAAT-HAP5 |
| AT1G54830 | CCAAT-HAP5 |
| AT1G56170 | CCAAT-HAP5 |
| AT3G12480 | CCAAT-HAP5 |
| AT3G22760 | CPP |
| AT4G14770 | CPP |
| AT1G47870 | E2F/DP |
| AT2G36010 | E2F/DP |
| AT3G01330 | E2F/DP |
| AT5G02470 | E2F/DP |
| AT5G03415 | E2F/DP |
| AT1G73730 | EIL |
| AT2G27050 | EIL |
| AT3G20770 | EIL |
| AT5G21120 | EIL |
| AT1G67710 | GARP-ARR-B |
| AT2G01760 | GARP-ARR-B |
| AT2G25180 | GARP-ARR-B |
| AT3G16857 | GARP-ARR-B |
| AT3G62670 | GARP-ARR-B |
| AT4G31920 | GARP-ARR-B |
| AT3G24120 | GARP-G2-like |
| AT4G17695 | GARP-G2-like |
| AT4G18020 | GARP-G2-like |
| AT5G16560 | GARP-G2-like |
| AT5G42630 | GARP-G2-like |
| AT5G58080 | GARP-G2-like |
| AT1G44810 | GeBP |
| AT1G61730 | GeBP |
| AT2G25650 | GeBP |
| AT4G25210 | GeBP |
| AT5G28040 | GeBP |
| AT1G07520 | GRAS |
| AT1G07530 | GRAS |
| AT1G14920 | GRAS |
| AT1G21450 | GRAS |
| AT1G50420 | GRAS |
| AT1G50600 | GRAS |
| AT1G55580 | GRAS |
| AT1G63100 | GRAS |
| AT1G66350 | GRAS |
| AT2G01570 | GRAS |
| AT2G04890 | GRAS |
| AT2G29060 | GRAS |
| AT2G45160 | GRAS |
| AT3G03450 | GRAS |
| AT3G46600 | GRAS |
| AT3G54220 | GRAS |
| AT3G60630 | GRAS |
| AT4G00150 | GRAS |
| AT4G17230 | GRAS |
| AT4G36710 | GRAS |
| AT4G37650 | GRAS |
| AT5G41920 | GRAS |
| AT5G48150 | GRAS |
| AT5G52510 | GRAS |

| | |
|---|---|
| AT5G59450 | GRAS |
| AT2G22840 | GRF |
| AT2G36400 | GRF |
| AT4G37740 | GRF |
| AT1G19700 | HB |
| AT1G20700 | HB |
| AT1G23380 | HB |
| AT1G26960 | HB |
| AT1G27050 | HB |
| AT1G30490 | HB |
| AT1G46480 | HB |
| AT1G52150 | HB |
| AT1G62360 | HB |
| AT1G62990 | HB |
| AT1G69780 | HB |
| AT1G70510 | HB |
| AT1G75410 | HB |
| AT1G79840 | HB |
| AT2G16400 | HB |
| AT2G17950 | HB |
| AT2G22430 | HB |
| AT2G22800 | HB |
| AT2G23760 | HB |
| AT2G28610 | HB |
| AT2G33880 | HB |
| AT2G34710 | HB |
| AT2G35940 | HB |
| AT2G46680 | HB |
| AT3G01220 | HB |
| AT3G19510 | HB |
| AT3G61890 | HB |
| AT4G00730 | HB |
| AT4G08150 | HB |
| AT4G16780 | HB |
| AT4G17460 | HB |
| AT4G32040 | HB |
| AT4G32880 | HB |
| AT4G34610 | HB |
| AT4G35550 | HB |
| AT4G36740 | HB |
| AT4G36870 | HB |
| AT4G36870 | HB |
| AT4G37790 | HB |
| AT4G40060 | HB |
| AT5G03790 | HB |
| AT5G06710 | HB |
| AT5G11060 | HB |
| AT5G25220 | HB |
| AT5G44180 | HB |
| AT5G47370 | HB |
| AT5G53980 | HB |
| AT5G59340 | HB |
| AT5G60690 | HB |
| AT5G65310 | HB |
| AT1G32330 | HSF |
| AT1G67970 | HSF |
| AT1G77570 | HSF |
| AT2G26150 | HSF |
| AT3G02990 | HSF |
| AT3G22830 | HSF |
| AT3G24520 | HSF |
| AT3G51910 | HSF |
| AT3G63350 | HSF |
| AT4G11660 | HSF |
| AT4G13980 | HSF |
| AT4G18880 | HSF |
| AT5G45710 | HSF |
| AT5G62020 | HSF |

| | |
|---|---|
| AT1G30810 | JUMONJI |
| AT1G63490 | JUMONJI |
| AT2G38950 | JUMONJI |
| AT4G20400 | JUMONJI |
| AT5G61850 | LFY |
| AT2G32700 | LUG |
| AT1G17310 | MADS |
| AT1G24260 | MADS |
| AT1G26310 | MADS |
| AT1G69120 | MADS |
| AT1G71692 | MADS |
| AT2G14210 | MADS |
| AT2G22540 | MADS |
| AT2G42830 | MADS |
| AT2G45660 | MADS |
| AT3G54340 | MADS |
| AT3G57230 | MADS |
| AT3G57390 | MADS |
| AT3G58780 | MADS |
| AT4G18960 | MADS |
| AT4G24540 | MADS |
| AT4G37940 | MADS |
| AT5G10140 | MADS |
| AT5G13790 | MADS |
| AT5G15800 | MADS |
| AT5G20240 | MADS |
| AT5G48670 | MADS |
| AT5G60910 | MADS |
| AT1G06180 | MYB |
| AT1G09540 | MYB |
| AT1G09770 | MYB |
| AT1G14350 | MYB |
| AT1G16490 | MYB |
| AT1G17950 | MYB |
| AT1G18570 | MYB |
| AT1G22640 | MYB |
| AT1G34670 | MYB |
| AT1G48000 | MYB |
| AT1G63910 | MYB |
| AT1G66230 | MYB |
| AT1G73410 | MYB |
| AT1G74080 | MYB |
| AT1G74430 | MYB |
| AT1G79180 | MYB |
| AT2G16720 | MYB |
| AT2G23290 | MYB |
| AT2G31180 | MYB |
| AT2G37630 | MYB |
| AT2G38090 | MYB |
| AT2G47190 | MYB |
| AT2G47460 | MYB |
| AT3G02940 | MYB |
| AT3G06490 | MYB |
| AT3G08500 | MYB |
| AT3G09370 | MYB |
| AT3G11280 | MYB |
| AT3G11440 | MYB |
| AT3G23250 | MYB |
| AT3G24310 | MYB |
| AT3G28910 | MYB |
| AT3G46130 | MYB |
| AT3G48920 | MYB |
| AT3G49690 | MYB |
| AT3G50060 | MYB |
| AT3G55730 | MYB |
| AT3G61250 | MYB |
| AT3G62610 | MYB |
| AT4G01680 | MYB |

| | |
|---|---|
| AT4G05100 | MYB |
| AT4G09460 | MYB |
| AT4G12350 | MYB |
| AT4G18770 | MYB |
| AT4G22680 | MYB |
| AT4G25560 | MYB |
| AT4G28110 | MYB |
| AT4G34990 | MYB |
| AT4G37260 | MYB |
| AT4G38620 | MYB |
| AT5G01200 | MYB |
| AT5G04760 | MYB |
| AT5G05790 | MYB |
| AT5G06100 | MYB |
| AT5G07690 | MYB |
| AT5G07700 | MYB |
| AT5G08520 | MYB |
| AT5G10280 | MYB |
| AT5G11510 | MYB |
| AT5G12870 | MYB |
| AT5G14340 | MYB |
| AT5G14750 | MYB |
| AT5G16600 | MYB |
| AT5G16770 | MYB |
| AT5G17800 | MYB |
| AT5G26660 | MYB |
| AT5G40330 | MYB |
| AT5G49620 | MYB |
| AT5G52260 | MYB |
| AT5G54230 | MYB |
| AT5G57620 | MYB |
| AT5G58900 | MYB |
| AT5G59780 | MYB |
| AT5G60890 | MYB |
| AT5G61420 | MYB |
| AT5G65790 | MYB |
| AT5G67300 | MYB |
| AT1G01060 | MYB-related |
| AT1G01380 | MYB-related |
| AT1G17520 | MYB-related |
| AT1G18330 | MYB-related |
| AT1G19000 | MYB-related |
| AT1G49950 | MYB-related |
| AT1G58220 | MYB-related |
| AT1G70000 | MYB-related |
| AT1G74840 | MYB-related |
| AT2G13960 | MYB-related |
| AT2G46830 | MYB-related |
| AT3G16350 | MYB-related |
| AT4G39160 | MYB-related |
| AT5G02840 | MYB-related |
| AT5G37260 | MYB-related |
| AT5G47390 | MYB-related |
| AT5G52660 | MYB-related |
| AT5G67580 | MYB-related |
| AT1G01010 | NAC |
| AT1G01720 | NAC |
| AT1G02220 | NAC |
| AT1G12260 | NAC |
| AT1G25580 | NAC |
| AT1G28470 | NAC |
| AT1G32870 | NAC |
| AT1G33060 | NAC |
| AT1G34180 | NAC |
| AT1G34190 | NAC |
| AT1G52880 | NAC |
| AT1G52890 | NAC |
| AT1G54330 | NAC |

| | |
|---|---|
| AT1G56010 | NAC |
| AT1G62700 | NAC |
| AT1G69490 | NAC |
| AT1G71930 | NAC |
| AT1G76420 | NAC |
| AT1G77450 | NAC |
| AT2G18060 | NAC |
| AT2G27300 | NAC |
| AT2G33480 | NAC |
| AT3G03200 | NAC |
| AT3G04070 | NAC |
| AT3G10500 | NAC |
| AT3G15170 | NAC |
| AT3G15500 | NAC |
| AT3G15510 | NAC |
| AT3G17730 | NAC |
| AT3G29035 | NAC |
| AT3G49530 | NAC |
| AT4G01550 | NAC |
| AT4G27410 | NAC |
| AT4G28500 | NAC |
| AT4G29230 | NAC |
| AT4G36160 | NAC |
| AT5G04410 | NAC |
| AT5G07680 | NAC |
| AT5G09330 | NAC |
| AT5G13180 | NAC |
| AT5G14000 | NAC |
| AT5G18270 | NAC |
| AT5G22290 | NAC |
| AT5G24590 | NAC |
| AT5G46590 | NAC |
| AT5G50820 | NAC |
| AT5G53950 | NAC |
| AT5G62380 | NAC |
| AT5G63790 | NAC |
| AT5G64530 | NAC |
| AT5G66300 | NAC |
| AT1G24030 | nFT |
| AT1G32130 | nFT |
| AT1G67910 | nFT |
| AT1G72060 | nFT |
| AT2G22900 | nFT |
| AT2G40670 | nFT |
| AT3G14000 | nFT |
| AT4G00760 | nFT |
| AT4G29410 | nFT |
| AT5G54930 | nFT |
| AT5G64220 | nFT |
| AT1G18790 | Nin-like |
| AT1G20640 | Nin-like |
| AT1G64530 | Nin-like |
| AT1G76350 | Nin-like |
| AT2G17150 | Nin-like |
| AT2G43500 | Nin-like |
| AT4G24020 | Nin-like |
| AT5G53040 | Nin-like |
| AT1G05380 | PHD |
| AT2G36720 | PHD |
| AT1G53160 | SBP |
| AT1G69170 | SBP |
| AT1G76580 | SBP |
| AT2G33810 | SBP |
| AT5G18830 | SBP |
| AT5G43270 | SBP |
| AT5G50670 | SBP |
| AT5G66350 | SRS |
| AT1G35560 | TCP |

| | |
|---|---|
| AT1G58100 | TCP |
| AT2G31070 | TCP |
| AT2G45680 | TCP |
| AT3G27010 | TCP |
| AT3G47620 | TCP |
| AT5G51910 | TCP |
| AT1G13450 | Trihelix |
| AT1G33240 | Trihelix |
| AT1G54060 | Trihelix |
| AT1G76880 | Trihelix |
| AT1G76890 | Trihelix |
| AT2G44730 | Trihelix |
| AT3G10040 | Trihelix |
| AT3G14180 | Trihelix |
| AT3G24490 | Trihelix |
| AT3G54390 | Trihelix |
| AT5G01380 | Trihelix |
| AT5G28300 | Trihelix |
| AT1G25280 | TUB |
| AT1G47270 | TUB |
| AT1G53320 | TUB |
| AT1G76900 | TUB |
| AT2G18280 | TUB |
| AT2G47900 | TUB |
| AT3G06380 | TUB |
| AT5G18680 | TUB |
| AT4G28190 | ULT |
| AT1G13960 | WRKY |
| AT1G29280 | WRKY |
| AT1G62300 | WRKY |
| AT1G68150 | WRKY |
| AT1G69310 | WRKY |
| AT1G69810 | WRKY |
| AT1G80840 | WRKY |
| AT2G04880 | WRKY |
| AT2G23320 | WRKY |
| AT2G24570 | WRKY |
| AT2G30250 | WRKY |
| AT2G30590 | WRKY |
| AT2G38470 | WRKY |
| AT2G40740 | WRKY |
| AT2G46400 | WRKY |
| AT2G47260 | WRKY |
| AT3G01970 | WRKY |
| AT3G56400 | WRKY |
| AT3G58710 | WRKY |
| AT4G01250 | WRKY |
| AT4G23810 | WRKY |
| AT4G24240 | WRKY |
| AT4G26640 | WRKY |
| AT4G30935 | WRKY |
| AT4G31550 | WRKY |
| AT5G13080 | WRKY |
| AT5G15130 | WRKY |
| AT5G22570 | WRKY |
| AT5G49520 | WRKY |
| AT5G56270 | WRKY |
| AT1G14440 | ZF-HD |
| AT1G69600 | ZF-HD |
| AT1G74660 | ZF-HD |
| AT1G75240 | ZF-HD |
| AT2G02540 | ZF-HD |
| AT3G28920 | ZF-HD |
| AT5G15210 | ZF-HD |
| AT5G39760 | ZF-HD |

**Supplemental Table2. List of domesticated parts and T.U in Goldenbraid**

| | **Domestication** | |
|---|---|---|
| **GBparts** | **Vector** | **Reference** |
| pCUC3 | pUPD2 | |
| *CUC3* CDS | pUPD2 | |
| pCUC3-6m | pUPD2 | |
| *CUC3-1m* CDS | pUPD2 | |
| Terminator 35S | pUPD2 | Sarrion-Perdigones et al., 2013 |
| GFP | pUPD2 | Sarrion-Perdigones et al., 2013 |
| mCherry | pUPD2 | Sarrion-Perdigones et al., 2013 |
| N7 | pUPD2 | |
| | **First level of T.U** | |
| **T.U** | **Vector** | **Reference** |
| pCUC3:CUC3:t35s | pDGB3_α1 | |
| pCUC3-6m:CUC3-1m:t35s | pDGB3_α1 | |
| pCUC3:mCherry-N7:t35s | pDGB3_α1 | |
| pCUC3-6m:GFP-N7:t35s | pDGB3_α1 | |
| pnos:hygro:tnos | pDGB3_α2 | Sarrion-Perdigones et al., 2013 |
| pCMV:DSRed:tnos | pDGB3_α2 | Morineau et al., 2017 |
| | **Final level of T.U** | |
| **T.U** | **Vector** | **Reference** |
| pCUC3:CUC3:t35s pCMV:DSRed:tnos | pDGB3_Ω1 | |
| pCUC3-6m:CUC3-1m:t35s pCMV:DSRed:tnos | pDGB3_Ω1 | |
| pCUC3:mCherry-N7:t35s pnos:hygro:tnos | pDGB3_Ω1 | |
| PCUC3-6m:GFP-N7:t35s pnos:hygro:tnos | pDGB3_Ω1 | |


Morineau, C., Bellec, Y., Tellier, F., Gissot, L., Kelemen, Z., Nogué, F., and Faure, J.D. (2017). Selective gene dosage by CRISPR-Cas9 genome editing in hexaploid Camelina sativa. Plant Biotechnol. J. 15, 729–739.

Sarrion-Perdigones, A., Vazquez-Vilar, M., Palací, J., Castelijns, B., Forment, J., Ziarsolo, P., Blanca, J., Granell, A., and Orzaez, D. (2013). Goldenbraid 2.0: A comprehensive DNA assembly framework for plant synthetic biology. Plant Physiol. 162, 1618–1631.

**Supplemental Table 4. Acquisition parameters for confocal imaging**

| Fluorescent Molecule | Excitation wavelength (nm) | Detection wavelength (nm) |
|---|---|---|
| Calcofluor | **405** | **410-450** |
| pCUC2:erRFP | **561** | **570-635** |
| pCUC3:erCFP | **458** | **460-512** |
| pWUS:VENUS-NLS | **514** | **518-543** |
| pCLV3:mCHERRY-NLS | **561** | **590-650** |
| pSOD7:GFP | **488** | **518-554** |
| pDPA4:GFP | **488** | **518-554** |
| Vector Blue (in situ) | **633** | **712-796** |
